# AP-1 specifies developmental versus fibrotic extracellular matrix transcriptional programs in the lung

**DOI:** 10.64898/2026.09.10.750752

**Authors:** Alyssa M. Kaiser, Angela Luo, Kathyayini Sivasubramanian, Georgina Dueñas, Baby Martin-McNulty, Wendy Cedron-Craft, John M. Patino, Andy J. Chang, Johannes Riegler, J. Graham Ruby

## Abstract

Lung function requires an elastic extracellular matrix (ECM). However, the adult lung does not regenerate the elastic architecture established during development, and deposition of fibrous ECM characterizes many lung diseases. We generated a multimodal, single-nuclei atlas across development, homeostasis, aging, and fibrosis to identify lung fibroblast populations, transcriptional programs, and regulatory logic governing these divergent matrix outcomes. We discovered differential AP-1 transcription factor activity orchestrates distinct ECM programs via preferential binding to TPA-responsive elements (TRE) in fibrosis and cAMP-responsive elements (CRE) in development. Antagonizing AP-1 TRE-signaling in lung fibroblasts repressed the fibrotic program and re-engaged a developmental, elastogenic state, and MEK inhibition produced similar transcriptional phenotypes. Pathological fibroblasts in interstitial lung diseases upregulated TRE motif activity, and expression of downstream fibrosis signatures positively correlated with disease severity. These results uncover AP-1 as a critical signaling hub governing lung fibroblast ECM deposition that can potentially be exploited to improve disease outcomes.

## Introduction

The extracellular matrix (ECM) is a complex, tissue-specific network of macromolecules including collagens, elastic fibers, glycoproteins, and proteoglycans^1,2^. The ECM provides a specialized architecture and signaling scaffold that governs the mechanical properties and proper function of diverse tissues, including the lung. In a healthy lung, the ECM provides the elastic recoil and distensibility required for ventilation and the structural support for alveoli to facilitate gas exchange^1,2^. Lung ECM is deposited throughout development, and its composition and structure depends on the anatomical location and developmental stage^1,3–5^. A key timepoint for ECM synthesis is alveolarization, when elastogenesis occurs to provide support for alveolar morphogenesis and long-term mechanical compliance^1,3–7^. The adult lung has a poor capacity for elastic tissue repair and regeneration, and elastic recoil is progressively lost with age, diminishing the lung’s capacity for efficient ventilation^8,9^. The inability to maintain and regenerate elastic ECM compromises lung function and contributes to a progressive decline in respiratory function over time^8–10^.

Failure to maintain the ECM architecture also underlies numerous lung diseases, such as interstitial lung disease (ILD) and chronic obstructive pulmonary disease (COPD)^1,11^. Idiopathic pulmonary fibrosis (IPF) is a progressive and fatal type of ILD characterized by the accumulation of fibrosis within the lung interstitium^12,13^. The deposition of stiff, collagen-dense ECM replaces native elastic architecture with non-compliant scar tissue, leading to a decrease in lung function and respiratory failure^12–14^. Despite the high mortality associated with the disease, current IPF therapies only slow the rate of functional decline rather than reverse established fibrosis or restore lung elasticity^12,15,16^. Restoring ECM homeostasis in IPF would require both suppressing fibrosis and reactivating elastogenic programs. Redirecting fibroblasts from synthesizing fibrotic towards tissue-appropriate ECM could therefore reverse fibrosis while rehabilitating the mechanical properties of the lung. However, the cell states that drive these ECM programs and the regulatory mechanisms that underlie them remain poorly understood. Elucidating mechanisms regulating ECM production across different contexts is essential to understand how fibroblast states affect matrisome deposition and to devise therapies capable of achieving functional lung repair.

AP-1 (activator protein 1) is a dimeric transcription factor complex, composed of two monomers from the JUN, FOS, ATF, CREB, and MAF protein families^17,18^. AP-1 family members contain a bZIP domain that enables their dimerization and docking to DNA as well as a transactivation domain to affect downstream gene expression^17,18^. The specific combination of dimerizing family members dictates motif preference, which include TPA-responsive elements (TRE), CRE (cAMP-responsive elements), and MAF recognition elements (MARE), each associated with specific genes and downstream transcriptional networks^17,19,20^. AP-1 activity has been previously associated with tissue injury, remodeling, and fibrosis; however, how AP-1 regulates these processes is not understood^21–23^.

Here, we generated a single-cell, multimodal dataset of the lung across distinct ECM contexts - in development, homeostasis, aging, and fibrosis. Using gene expression (RNA-seq) and chromatin accessibility (ATAC-seq) modalities measured in the same nuclei, we identified myofibroblast populations and transcriptional programs that drive ECM deposition in development and disease. We discovered that differential AP-1 transcription factor activity underlies these divergent ECM gene expression programs: AP-1 preferentially binds TRE motifs in fibrotic, activated myofibroblasts and CRE motifs in developmental myofibroblasts. Inhibiting TRE motif binding *in vitro* repressed fibrotic ECM gene expression programs and induced a developmental, elastic ECM program, and small molecule inhibitor experiments revealed mitogen activated protein kinase (MAPK) signaling as a driver of fibrotic AP-1 activity. Importantly, TRE motif activity was elevated in ILDs, including IPF, and elevated expression of the TRE-associated fibrosis signature in IPF correlated with more severe disease. Our findings reveal a central pillar of regulatory logic in lung fibroblasts and bring to light a potential new therapeutic strategy to inhibit fibrosis and promote functional restoration of the lung ECM.

## Results

### Joint transcriptome and chromatin profiling resolves lung fibroblast states in development and disease

We designed an experiment to capture pathological and physiological ECM states to delineate transcriptional programs and regulatory drivers of distinct fibroblast phenotypes. Single-nuclei multiomics RNA- and ATAC-sequencing was performed on mouse lungs across four distinct biological states: development, homeostasis, aging, and fibrosis (Fig. 1a). This design encompassed a continuum of ECM programs ranging from physiological synthesis of healthy ECM to dysregulation in age and disease and allowed cell populations, transcriptional programs, and regulatory elements to be compared across contexts. Fibrosis was modeled using bleomycin to induce acute lung injury^24,25^. A single dose of saline or bleomycin was administered intratracheally to 16-week old mice and lungs were collected 14 days thereafter (Fig. 1a). Bleomycin treatment resulted in weight loss over time, and micro-computed tomography (μCT) confirmed an increase in lung fibrosis/inflammation in all bleomycin-treated mice, as quantified by elevated high signal volume fraction (Supp. Fig. 1a-f)^26^. To study age-related changes to ECM programs, lungs were collected from postnatal day 7 (P7), adult (16-week old), and aged (90-week old) mice (Fig. 1a). μCT of the 16- and 90-week old animals revealed elevated expiratory lung volume and high signal intensity volume fraction in aged animals, suggesting an increase in inflammation and/or fibrosis (Supp. Fig. 1g-j)^26^.

**Figure 1:**
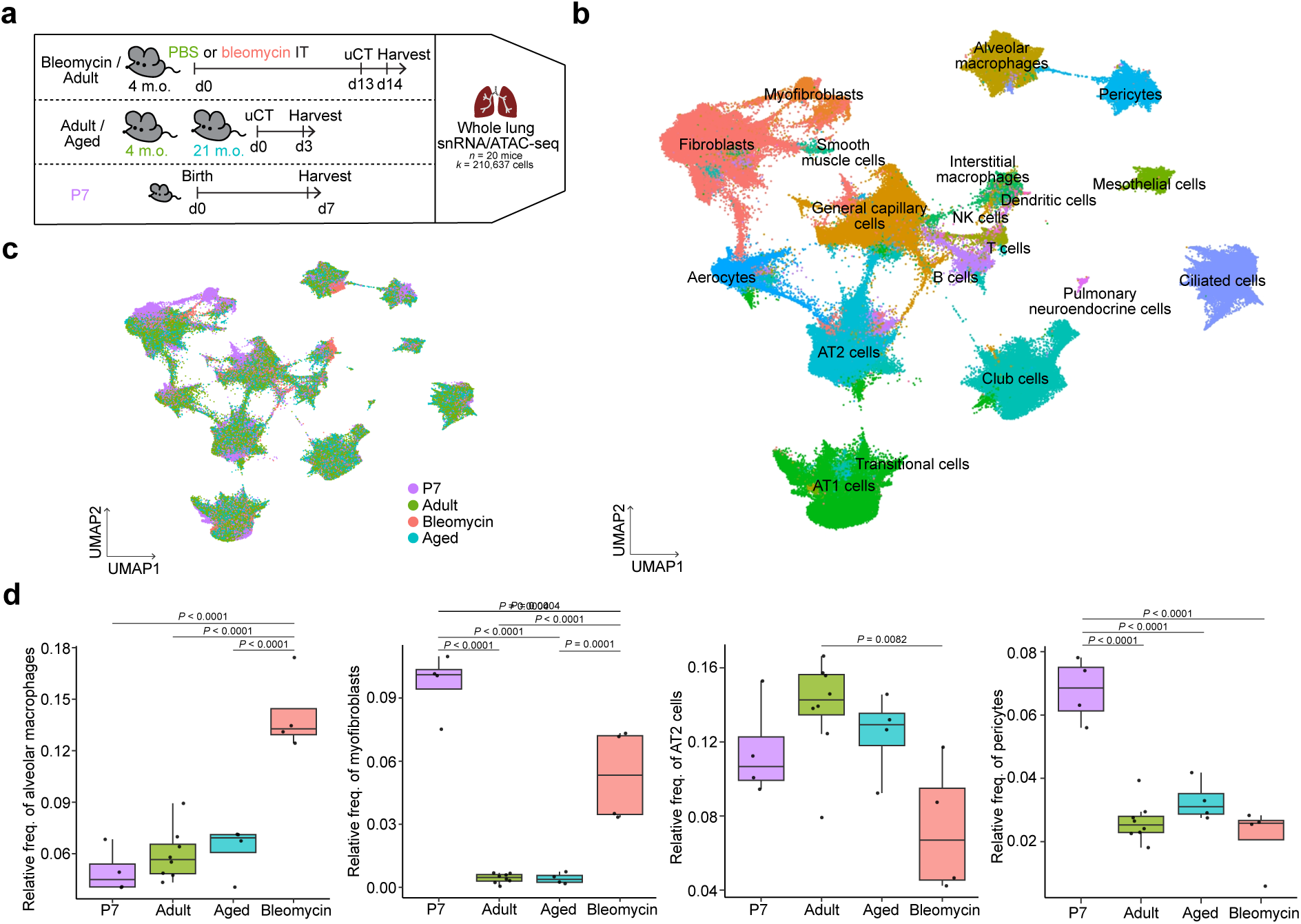
Multiomics analysis reveals cell population dynamics with age and disease. (a) Schematic for the multiomics experiment. Lungs were harvested from 4 month old (m.o.) mice treated intratracheally (IT) with either saline (*N* = 4) or bleomycin (*N* = 4) 14 days prior, untreated 4 m.o. mice (*N* = 4), untreated 21 m.o. mice (*N* = 4), and postnatal day 7 mice (*N* = 4) and snap frozen. Combined single nuclei RNA- and ATAC-seq was then performed on all frozen samples. uCT, micro-computed tomography. (b) UMAP projection of all cells after quality control filtering colored by cluster. (c) UMAP plot from (b) colored by condition. P7, postnatal day 7. (d) Box plots of proportions of various cell types across conditions in the whole dataset. Box plots display the median and interquartile range (IQR, 25th–75th percentiles), whiskers extend to the furthest data point within 1.5 x IQR of the lower and upper quartiles. *P* values were calculated by ordinary one-way ANOVA with Tukey’s multiple comparisons test (d).

To visualize the cellular architecture across conditions, the transcriptional and epigenetic modalities were jointly integrated into a unified multimodal uniform manifold approximation and projection (UMAP) embedding (Fig. 1b, Supp. Fig. 2a-b). The epithelial, mesenchymal, immune, and endothelial compartments were represented across conditions (Fig. 1b-c, Supp. Fig. 2c-f). Given the similarity between saline-treated and untreated 16-week old mice, these conditions were combined together as healthy, adult mice (Supp. Fig. 2e-f). While aged lungs exhibited minor deviations from the cell-type composition of healthy adult lungs, neonatal and bleomycin-injured lungs substantially diverged in the frequency of key cell types (Fig. 1c-d, Supp. Fig. 2f). For example, injured lungs showed an expansion of the immune and mesenchymal cell compartments, in particular alveolar macrophages and myofibroblasts (Fig. 1d, Supp. Fig. 2f).

To identify the transcriptional and epigenetic factors that drive ECM synthesis, we segmented out the fibroblast and myofibroblast populations (Fig. 2a). Annotation of fibroblast subtypes identified 8 main clusters across conditions. *Pdgfra*+ *Npnt*+ alveolar fibroblasts were the predominant population in adult and aged lungs, whereas *Dcn*+ *Pi16*+ adventitial fibroblasts and *Hhip*+ *Cdh4*+ airway ductal myofibroblasts constituted smaller populations (Fig. 2a-c, Supp. Fig. 3a)^27–30^. Additionally, healthy adult lungs contained matrix fibroblasts, a subtype of alveolar fibroblasts with elevated expression of ECM genes including *Col4a1* and *Fn1*, and this population was expanded in aged lungs (Fig. 2a-c, Supp. Fig. 3a-c)^30^. In bleomycin-treated lungs, alveolar fibroblasts remained the most abundant fibroblast type; however, their relative frequency was decreased due to the emergence of activated myofibroblasts. These pathologic fibroblasts have been previously described: they express markers including *Cthrc1* and *Spp1* as well as fibrotic ECM gene expression programs derived from studies of mouse radiation- and bleomycin-induced lung fibrosis (Fig. 2b-e, Supp. Fig. 3a-c, Supplementary Table 1)^27,31–34^.

**Figure 2:**
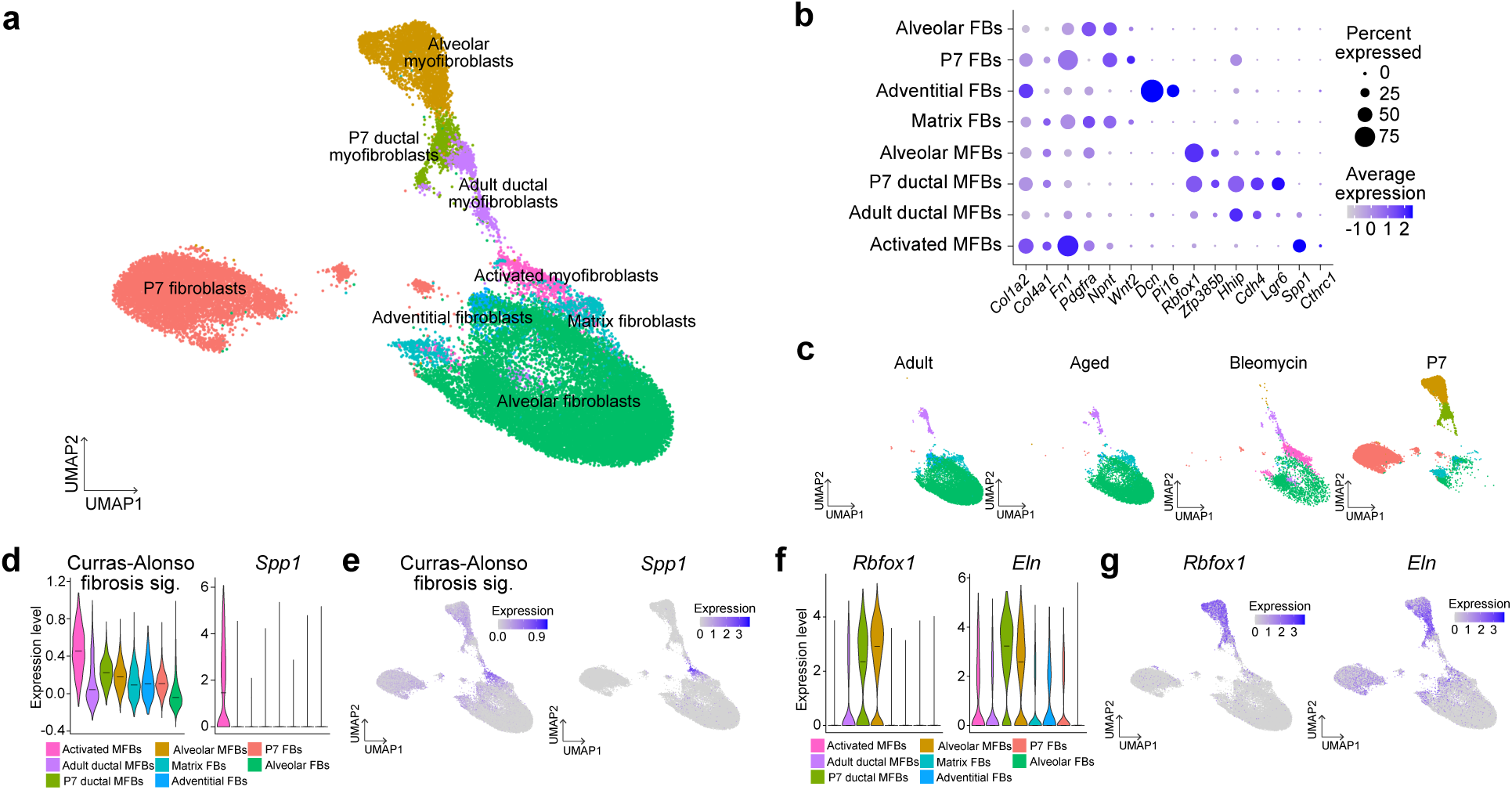
Age and disease context dictates the lung fibroblast landscape. (a) UMAP projection of all fibroblasts colored by cluster. (b) Dot plot of expression of fibroblast gene markers across all fibroblast populations. (c) UMAP plot from (a) split by condition. (d) Violin plots of expression of (left) a fibrosis gene signature derived from a mouse model of radiation-induced lung fibrosis^32^ and (right) *Spp1*, a marker of activated myofibroblasts, across fibroblast populations. FB, fibroblast; MFB, myofibroblast. (e) UMAP plot visualizations of (d). (f) (Left) Violin plots of developmental myofibroblast gene markers across fibroblast populations. (g) UMAP plot visualizations of (f). Violin plots show the median and the relative frequency of cells at a given expression value.

P7 lungs comprised developmentally-restricted states that were largely absent from the landscape of the adult lung (Fig. 2c, Supp. Fig. 3a). The largest population was *Wnt2*+ fibroblasts, a precursor to adult *Pdgfra*+ fibroblasts (Fig. 2a-c)^29,35^. The second largest population were developmental alveolar and ductal myofibroblasts, defined by co-expression of myofibroblast genes (including *Rbfox1* and *Zfp385b*) and alveolar (*Pdgfra*) and ductal (*Hhip*) specific markers (Fig. 2a-c, f-g, Supp. Fig. 3a-c)^4,29,36–41^. Matrix and adventitial fibroblasts were also minor populations present in neonatal lungs (Fig. 2a-c, Supp. Fig. 3a)^29,42^.

### Specialized ECM transcriptional programs distinguish fibrotic from developmental myofibroblasts

To determine if and how transcriptional regulation shapes the distinct ECM produced in disease and development, we defined the cell types and gene expression programs driving pathological and physiological ECM synthesis. We focused on activated and P7 myofibroblasts as the primary drivers of fibrotic and developmental ECM. Activated, pathogenic myofibroblasts are known to contribute to fibrosis and disease progression in mice and humans, and these myofibroblasts in our dataset robustly express fibrotic ECM programs, making them top candidates for driving adverse ECM outcomes (Fig. 2d, Supp. Fig. 3b-c)^27,31,32^. In development, P7 alveolar and ductal myofibroblasts expressed the highest levels of genes associated with elastic fiber formation, such as *Eln, Loxl2*, and *Fbn2*, consistent with their role in producing and remodeling elastic fibers during alveolarization (Fig. 2f-g, Supp. Fig. 3b-c)^3,4,7,29,36–41^. These expression profiles coupled with the known contribution of developmental myofibroblasts to elastic tissue formation implicate P7 myofibroblasts as primary drivers of physiological, elastic ECM synthesis at this timepoint.

We subsequently interrogated whether discrete ECM-related gene expression programs distinguished activated from developmental myofibroblasts (see Methods) (Fig. 3a-b, Supp. Fig. 3d-f). While some ECM-associated genes were similarly expressed in both populations, a unique set of ECM genes were differentially expressed between the two populations (Fig. 3a-b, Supp. Fig. 3d-f, Supplementary Table 1). These two programs, designated the fibrotic and developmental ECM gene expression programs, were specifically upregulated in the activated myofibroblast or the alveolar and ductal P7 myofibroblast populations, respectively (Fig. 3a-b, Supp. Fig. 3d).

**Figure 3:**
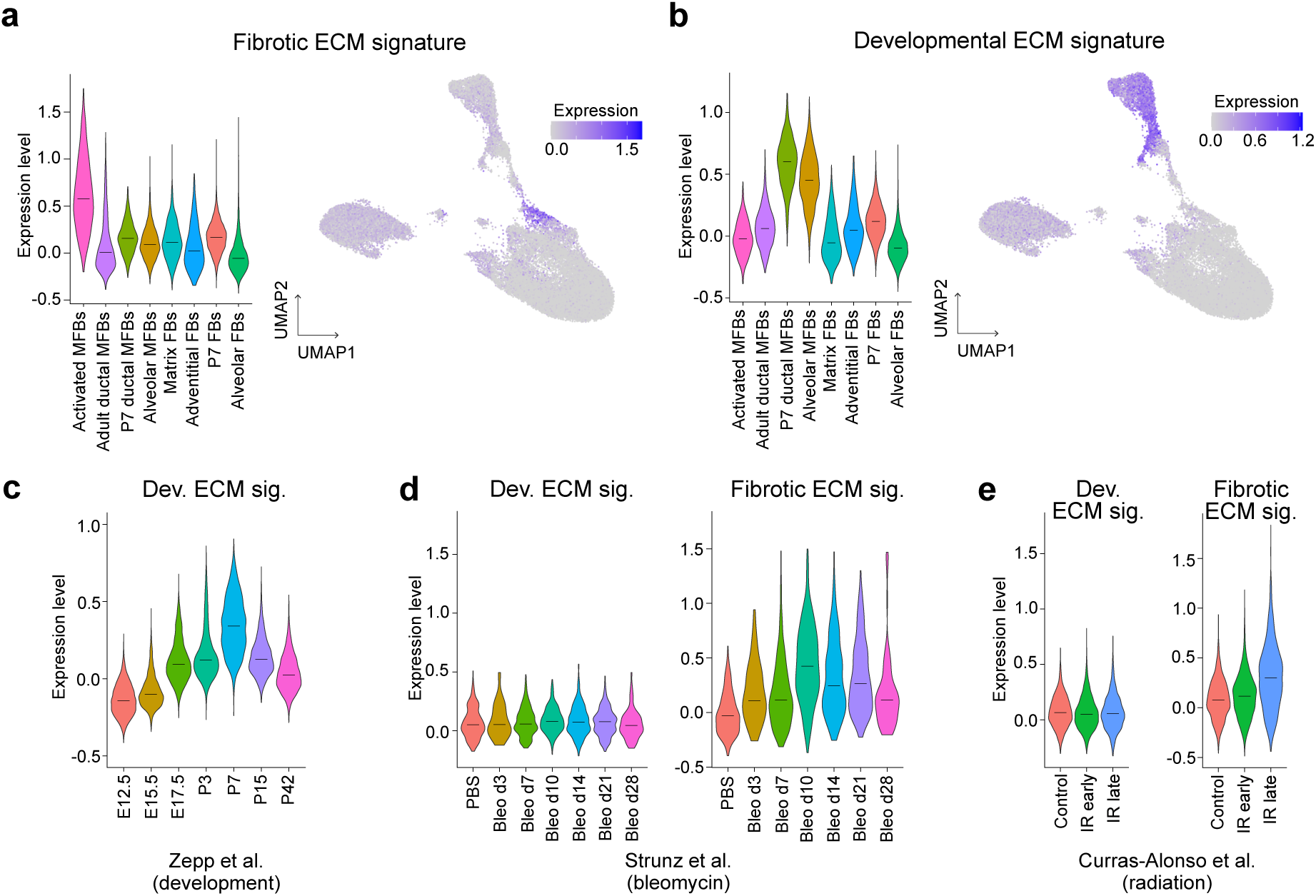
Myofibroblasts express distinct ECM transcriptional programs in disease and development. (a-b) (Left) Violin plots and (right) UMAP plots of expression of the (a) fibrotic or (b) developmental ECM program across fibroblast populations. (c) Violin plot of expression of the developmental ECM program in fibroblasts from an independent scRNA-seq dataset of mouse lung development^35^. Expression is shown in fibroblasts from embryonic day 12.5 (E12.5) to postnatal day 42 (P42) lungs. (d) Violin plots of expression of the (left) developmental and (right) fibrotic ECM programs in an independent scRNA-seq dataset of bleomycin-induced mouse lung injury^43^. Expression is shown in fibroblasts from control (PBS-treated) lungs or from bleomycin-treated lungs (3 to 28 days post-treatment). (e) Violin plots of expression of the (left) developmental and (right) fibrotic ECM programs in an independent scRNA-seq dataset of radiation-induced mouse lung injury^32^. Expression is shown in fibroblasts from control lungs, irradiation (IR) early lungs (17 Gy, 1-3 months post exposure), and IR late lungs (17 Gy, 4-5 months post exposure). Violin plots show the median and the relative frequency of cells at a given expression value.

The relevance and specificity of these expression programs were validated using independent datasets of mouse lung development and fibrosis. In two scRNA-seq experiments of mouse lung development, which comprised samples ranging from embryonic to adult lungs, the developmental ECM signature peaked in fibroblasts at postnatal day 7 and then decreased into adulthood (Fig. 3c, Supp. Fig. 3g)^29,35^. In contrast, the fibrotic gene expression signature remained stable in fibroblasts from late embryonic development (∼E19) to adult stage lungs (Supp. Fig. 3g-h). Meanwhile, the pathological ECM signature was induced in disease compared to control fibroblasts across three scRNA-seq experiments using bleomycin- or radiation-induced lung injury models (Fig. 3d-e, Supp. Fig. 3i-l)^27,32,43^. In bleomycin-exposed fibroblasts, expression of the fibrotic ECM signature peaked at 10 days post-treatment, while in radiation-treated fibroblasts the peak occurred at 4-5 months post-exposure (Fig. 3d-e, Supp. Fig. 3i-l). In contrast, the developmental ECM program was not induced in fibroblasts after either injury stimulus (Fig. 3d-e, Supp. Fig. 3i-l).

### Differential AP-1 transcription factor activity underlies distinct ECM transcriptional programs

To pinpoint transcription factors driving divergent myofibroblast identities and ECM programs, the ATAC-seq data was used to identify differentially accessible peaks across all fibroblast populations, including the developmental and activated myofibroblasts. Transcription factor motif discovery analysis (HOMER) was then performed to identify motifs enriched across these differential peaks and nominate transcription factors active in each population^44^. Interestingly, AP-1 transcription factor motifs were the top enriched motifs in both P7 and activated myofibroblasts; however, the specific AP-1 sequence differed between the two populations (Fig. 4a-c, Supplementary Tables 2, 3). The top motif enriched in developmental myofibroblasts was the cAMP response element (CRE motif), a palindromic eight-base pair sequence (5′ - <u>TGA</u>CG<u>TCA</u> - 3′) (Fig. 4a-b, Supplementary Table 2). Meanwhile, the top motif identified in activated myofibroblasts was the TPA response element (TRE motif), which differs in one base pair (5′ - <u>TGA</u>S<u>TCA</u> - 3′) (Fig. 4a, c, Supplementary Table 3). AP-1 motif preference is influenced by variables including dimer composition and cofactor interactions^17^. For example, JUN and FOS family proteins preferentially bind TRE motifs while ATF and CREB family members bias towards CRE motifs^17,45^.

**Figure 4:**
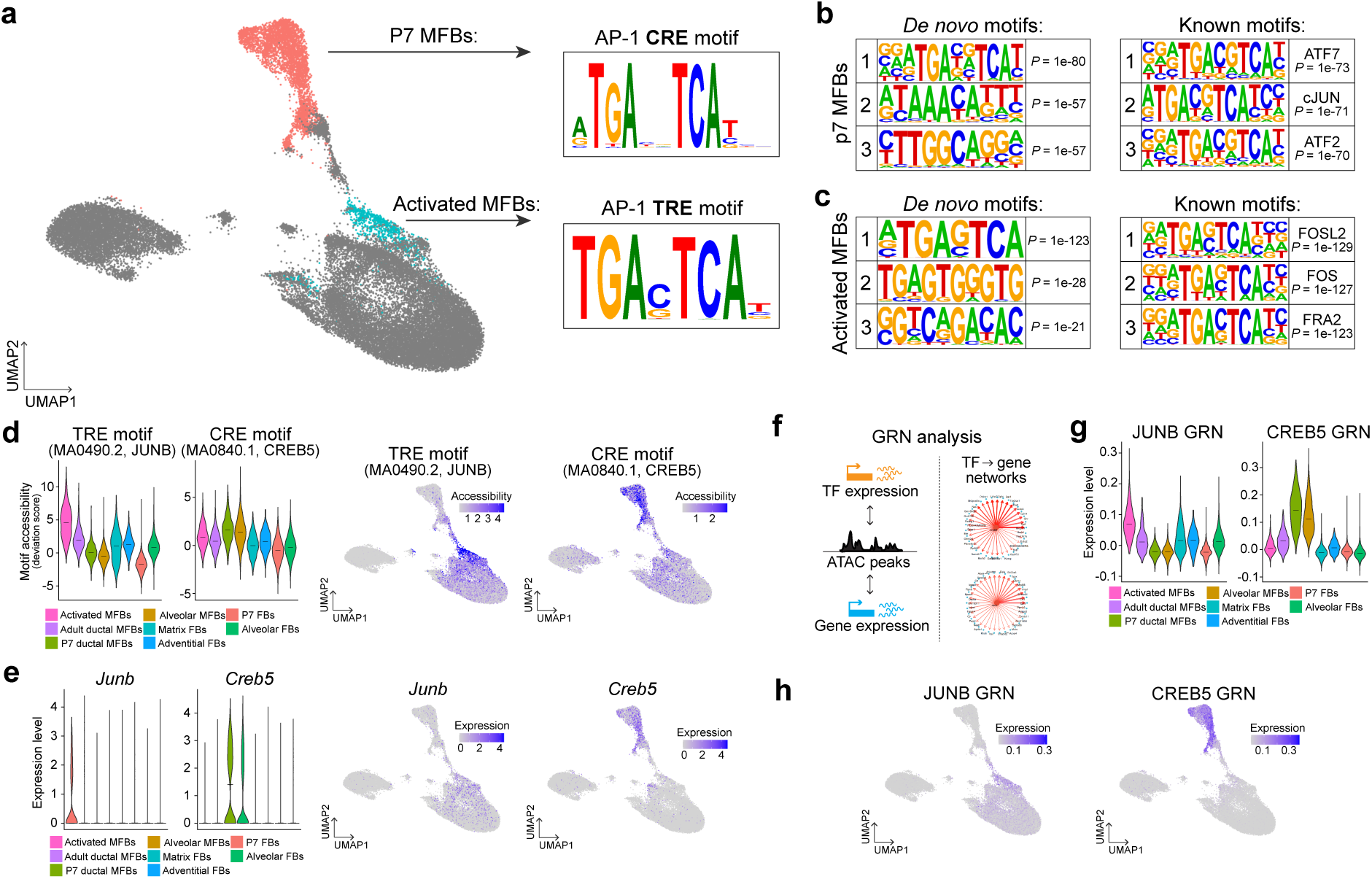
AP-1 transcription factor activity underlies myofibroblast states and ECM transcriptional programs. (a) Sequence logo plots of the top transcription factor motif enriched in differentially accessible peaks of developmental and activated myofibroblasts, identified by HOMER motif discovery analysis. (b-c) The top 3 (left) *de novo* and (right) known motifs by *P*-value enriched in differentially accessible peaks of (b) developmental or (c) activated myofibroblasts identified by HOMER. (d) (Left) Violin plots and (right) UMAP plots of motif accessibility (deviation score) of the JUNB and CREB5 motifs across fibroblast populations. (e) (Left) Violin plots and (right) UMAP plots of expression of *Junb* and *Creb5* across fibroblast populations. (f) (Left) Schematic visualizing the GRaNIE GRN pipeline^49^ and (right) representative network plots of identified transcription factor-gene networks. GRN, gene regulatory network. (g) Violin plots of expression of the identified JUNB and CREB5 GRNs identified across fibroblast populations. (h) UMAP plot visualizations of (g). Violin plots show the median and the relative frequency of cells at a given expression value. *P* values were calculated by cumulative hypergeometric test (b-c, HOMER).

We validated our motif discovery analysis using chromVAR to assess previously defined AP-1 family member motifs, which computes per-cell accessibility scores for annotated motifs^46^. TRE motifs associated with JUN and FOS family members were preferentially accessible in the activated myofibroblasts (Fig. 4d, Supp. Fig. 4a-b). Meanwhile, CRE motifs associated with CREB and ATF family members showed increased accessibility in the alveolar and ductal neonatal myofibroblasts (Fig. 4d, Supp. Fig. 4b-c). Importantly, across all cell populations identified in the whole lung dataset, accessibility of these AP-1 motifs was largely confined to these myofibroblast populations (Supp. Fig. 4d). The exception was elevated TRE accessibility in pre-alveolar type I transitional cells, a transient cell type that arises in lung injury (Supp. Fig. 4d)^43,47,48^. Thus, differential opening of TRE and CRE motifs broadly delineated pathologic and developmental myofibroblasts.

We next interrogated whether this distinct AP-1 regulatory landscape impacted ECM program expression. To identify transcription factors likely responsible for activating expression of the fibrotic and developmental programs, motif discovery analysis was restricted to differentially accessible regions between activated and developmental myofibroblasts linked to our ECM signature genes. The TRE and CRE motifs were significantly enriched in the accessible regions associated with the fibrotic and physiological ECM signature genes, respectively, suggesting AP-1 directly regulates these ECM programs (Supp. Fig. 4e, Supplementary Table 4).

To discern which family members contribute to AP-1 motif opening, we first examined the expression of TRE and CRE binders across contexts. *Junb* and *Fosl2* transcript levels were significantly elevated in activated myofibroblasts relative to developmental myofibroblasts, while *Creb5* and *Atf7* showed the inverse pattern (Fig. 4e, Supp. Fig. 4a-c). Next, gene regulatory networks (GRNs) were constructed using the multiomics data across all fibroblasts to computationally infer direct links between transcription factors, differentially accessible regulatory elements, and their target genes (Fig. 4f)^49^. Through this objective network construction, GRNs were discovered for the AP-1 family members CREB5, JUN, JUNB, FOS, and FOSL2 (Fig. 4f-h, Supp. Fig. 4f-g, Supplementary Tables 1, 5). Visualizing the expression of each AP-1 GRN revealed specific enrichment of the JUNB and FOSL2 networks in activated myofibroblasts and the CREB5 network in P7 myofibroblasts (Fig. 4g-h, Supp. Fig. 4f-g). Interestingly, CREB5 was predicted to directly regulate *Eln* expression, a gene essential for synthesizing elastic fibers (Supp. Fig. 4h, Supplementary Table 5). Altogether, these analyses highlight JUNB and FOSL2 as putative activators of the fibrotic ECM in activated myofibroblasts and CREB5 as a candidate to induce elastic ECM in developmental myofibroblasts.

### AP-1 directly modulates lung fibroblast ECM programs

Next, we directly tested if and how AP-1 regulates ECM programs in primary mouse lung fibroblasts *in vitro*, which in standard culture conditions transcriptionally most closely resembled activated myofibroblasts *in vivo* (Supp. Fig. 5a-b). To manipulate AP-1 TRE activity in these cells, we expressed a dominant negative AP-1 protein, AFOS, and performed RNA- and ATAC-sequencing (Fig. 5a, Supp. Fig. 5c-d)^50^. AFOS is a truncated and modified version of the FOS protein that lacks a functional DNA binding domain and contains an acidic extension that binds dimerization partners with a higher affinity than FOS, allowing it to sequester binding partners and inhibit target gene activation^50^. AFOS expression in mouse lung fibroblasts induced widespread transcriptional and epigenetic changes (Supp. Fig. 5e-f). AFOS expression significantly repressed accessibility of TRE motifs but not CRE motifs (Fig. 5b-c, Supp. Fig. 5g, Supplementary Table 6). TRE motifs repressed by AFOS expression were enriched at ATAC peak centers, implicating TRE motif occupancy in driving peak accessibility changes (Fig. 5d). A small group of TRE-containing peaks increased in accessibility in AFOS-expressing samples; however, this enrichment was not significant and the TRE motifs were dispersed throughout peak regions, suggesting they were not primary drivers of changing peak accessibility (Fig. 5b, e, Supp. Fig. 5g). In concordance with repressed TRE motif accessibility, AFOS expression robustly suppressed a previously described AP-1/JUN expression module, indicating the dominant negative inhibited TRE-mediated AP-1 signaling (Fig. 5f, Supp. Fig. 5h, Supplementary Table 1)^51^.

**Figure 5:**
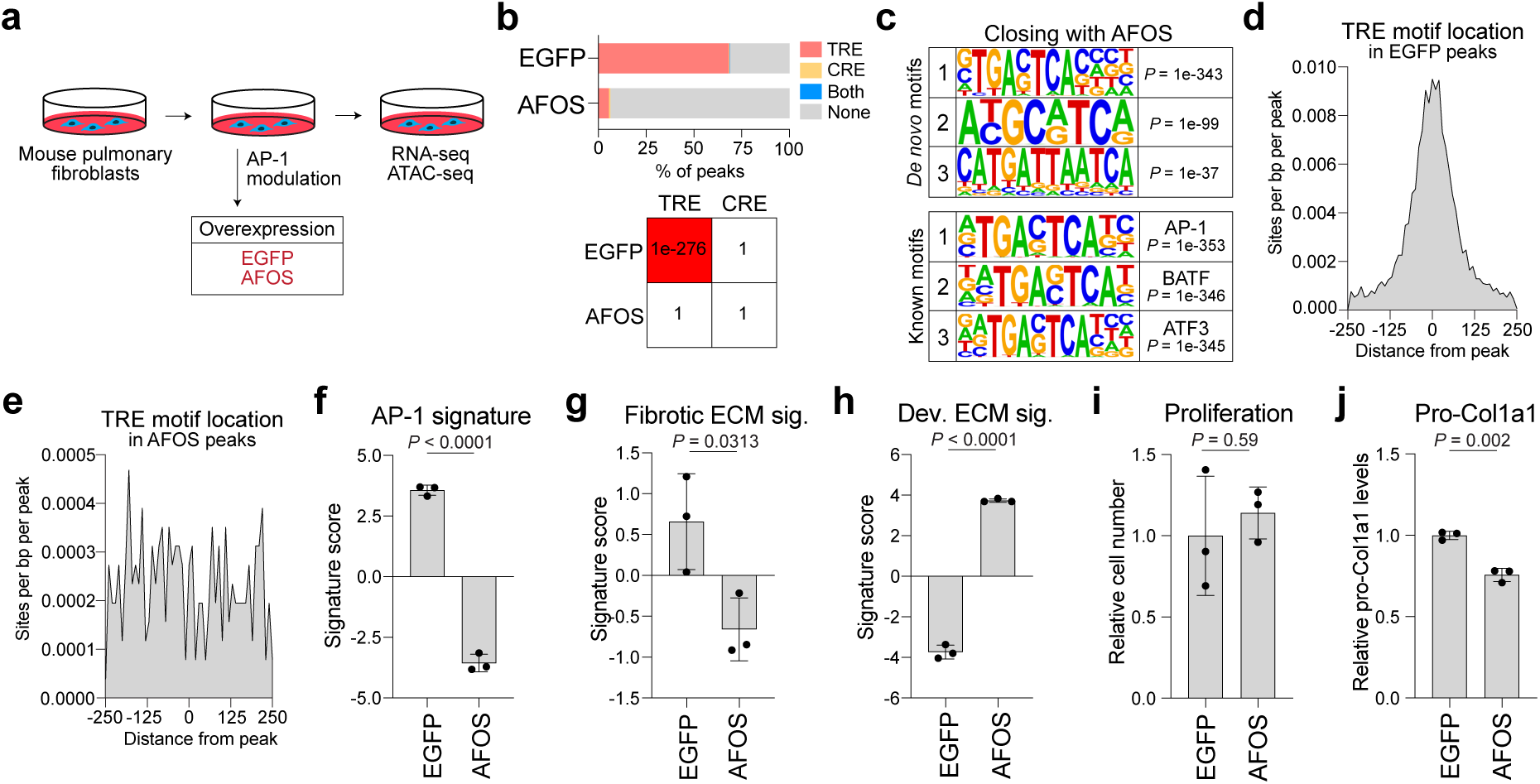
Inhibiting AP-1 TRE activity suppresses fibrotic and activates elastogenic ECM programs. (a) Schematic of the AFOS overexpression experiment. Mouse pulmonary fibroblasts (MPFs) were electroporated with expression constructs and harvested 24 hours later for RNA- and ATAC-seq (*N* = 3 per condition). (b) (Top) Bar graph of the proportion of differential peaks (*P*-adjusted < 0.05) increasing in accessibility in each condition that contain TRE and CRE motifs and (bottom) the *P*-value enrichment of these motifs across conditions. (c) The top 3 (top) *de novo* and (bottom) known motifs by *P*-value enriched in peaks significantly decreasing (*P*-adjusted < 0.05) in accessibility with AFOS expression in MPFs identified by HOMER. (d-e) Histograms of TRE motif frequency from peak centers of differentially accessible ATAC peaks in MPFs (*P* < 0.05) open in (d) control samples or (e) AFOS-expressing samples. (f) Bar graph of an AP-1 target gene signature^51^ composite expression score across conditions. (g-h) Bar graphs of the (g) fibrotic ECM signature or (h) developmental ECM signature composite expression scores across conditions. (i) Bar graph of the relative number of cells 24 hours after electroporation of EGFP or AFOS. (j) Bar graph of the relative level of pro-COL1A1 in MPFs 24 hours after electroporation of EGFP or AFOS. Bar graphs (f-j) show mean ± standard deviation. *P* values were calculated by cumulative hypergeometric test (c, HOMER) and by two-tailed Welch’s *t*-test (f-j).

AFOS expression induced widespread changes to ECM programs at the transcriptional and epigenetic levels. Differentially expressed genes were significantly enriched for ECM-related genes, including an induction of an elastic fiber synthesis program (Supp. Fig. 5i-j). Similarly, ECM-associated genes were overrepresented among genes linked to differentially accessible peaks (Supp. Fig. 5k). Given the extensive changes in ECM gene expression, we assessed how AFOS altered the fibrotic and developmental ECM programs. AFOS expression downregulated markers of activated myofibroblasts including *Spp1* and *Runx1* and the expression of the fibrotic ECM signature (Fig. 5g, Supp. Fig. 5l). Meanwhile, consistent with an upregulation of the elastic fiber synthesis program, AFOS expression robustly induced markers of P7 myofibroblasts such as *Eln* and the developmental ECM program expression (Fig. 5h, Supp. Fig. 5m). Thus, AFOS-mediated inhibition of the AP-1 TRE signaling axis drove a transcriptional switch in fibroblasts, repressing the pathological, fibrotic ECM program and inducing the developmental, elastic ECM program. In agreement with these transcriptional changes, levels of pro-COL1A1, a precursor for type I collagen, were significantly reduced in AFOS-expressing cells compared to control cells, while proliferation remained unchanged (Fig. 5i-j). Collectively, these results identify the AP-1 transcription factor complex as a master orchestrator of myofibroblast ECM states and a potential target for therapeutic modulation of fibrosis.

The contribution of individual AP-1 family members to these phenotypes was tested through systematic knockdown of JUN/FOS family members and overexpression of CRE binding AP-1 proteins CREB5 and ATF7, followed by RNA-sequencing (Supp. Fig. 6a). siRNA delivery decreased expression of all targeted transcripts with the exception of *Fos*, whose expression increased with delivery of any FOS family member siRNA (Supp. Fig. 6b-c). The impact of individual AP-1 member knockdown or overexpression on global transcription was minimal, with mild effects on AP-1 signaling as well as the developmental and fibrotic ECM programs (Supp. Fig. 6d-g). However, differentially expressed genes in several knockdown conditions, including *siJund* and *siFosl2*, were enriched for ECM-related genes (Supp. Fig. 5i). We also tested the combined knockdown of *Junb* or *Fosl1* with overexpression of *Creb5* or *Atf7*, which amplified the enrichment of ECM-related pathways within differentially expressed genes (Supp. Fig. 5i, Supp. Fig. 6a-b). Of all perturbations tested, combining *Junb* knockdown with *Creb5* overexpression elicited the most robust transcriptional response, including an induction of genes involved in elastic fiber synthesis (Supp. Fig. 5j, Supp. Fig. 6d). Thus, individual or dual manipulation of AP-1 family members was sufficient to elicit modest changes to ECM gene expression but inadequate to mimic phenotypes of AFOS expression, likely due to family member redundancy.

### MEK signaling regulates AP-1 activity in fibrotic fibroblasts

Diverse signaling inputs can regulate AP-1 activity at both the transcriptional and post-translational levels^17^. To identify upstream drivers of the AP-1 TRE-mediated ECM transcriptional response, we screened a small molecule inhibitor panel of known AP-1 regulators in fibroblasts (Fig. 6a)^17,52–57^. Inhibiting c-Jun N-terminal kinase (JNK) signaling, mitogen-activated protein kinase kinase (MEK) signaling, phosphoinositide 3-kinase (PI3K) signaling, transforming growth factor beta (TGFβ) signaling, JAK-STAT signaling, YAP signaling, and p38 MAPK signaling led to a downregulation of the AP-1/JUN gene expression signature, suggesting that these pathways activate AP-1 TRE signaling in lung fibroblasts (Fig. 6b, Supp. Fig. 6h-i). However, only MEK inhibition concurrently induced the developmental ECM program and repressed the fibrotic ECM program (Fig. 6c-d). MEK inhibition repressed expression of many TRE binding JUN/FOS family members including *Junb* and *Fosl1* and induced expression of CRE binding proteins like *Creb5* and *Atf7*, suggesting that transcriptional regulation of AP-1 family members at least partially explains MEK regulation of AP-1 signaling (Fig. 6e, Supp. Fig. 6j). Supporting the conclusion that MEK regulates these ECM programs through AP-1, the transcriptome-wide response of MEK inhibition was significantly correlated with that of AP-1 inhibition via AFOS across the top variable genes and ECM gene expression programs (Supp. Fig. 6k). These results suggested that MAPK signaling drives AP-1 signaling in pathological myofibroblasts *in vivo*. Indeed, MAPK/extracellular signal-regulated kinase (ERK) target genes were elevated in activated myofibroblasts relative to P7 myofibroblasts, whereas genes induced by MEK inhibition via Trametinib were upregulated in developmental myofibroblasts (Fig. 6f, Supp. Fig. 6l) (Supplementary Table 1)^58^. Altogether, these results lead to a model where the balance between AP-1 CRE and TRE activity determines whether elastic, physiologically appropriate or pathological, fibrotic ECM is synthesized, with differential MAPK/ERK activity tuning this activity (Fig. 6g).

**Figure 6:**
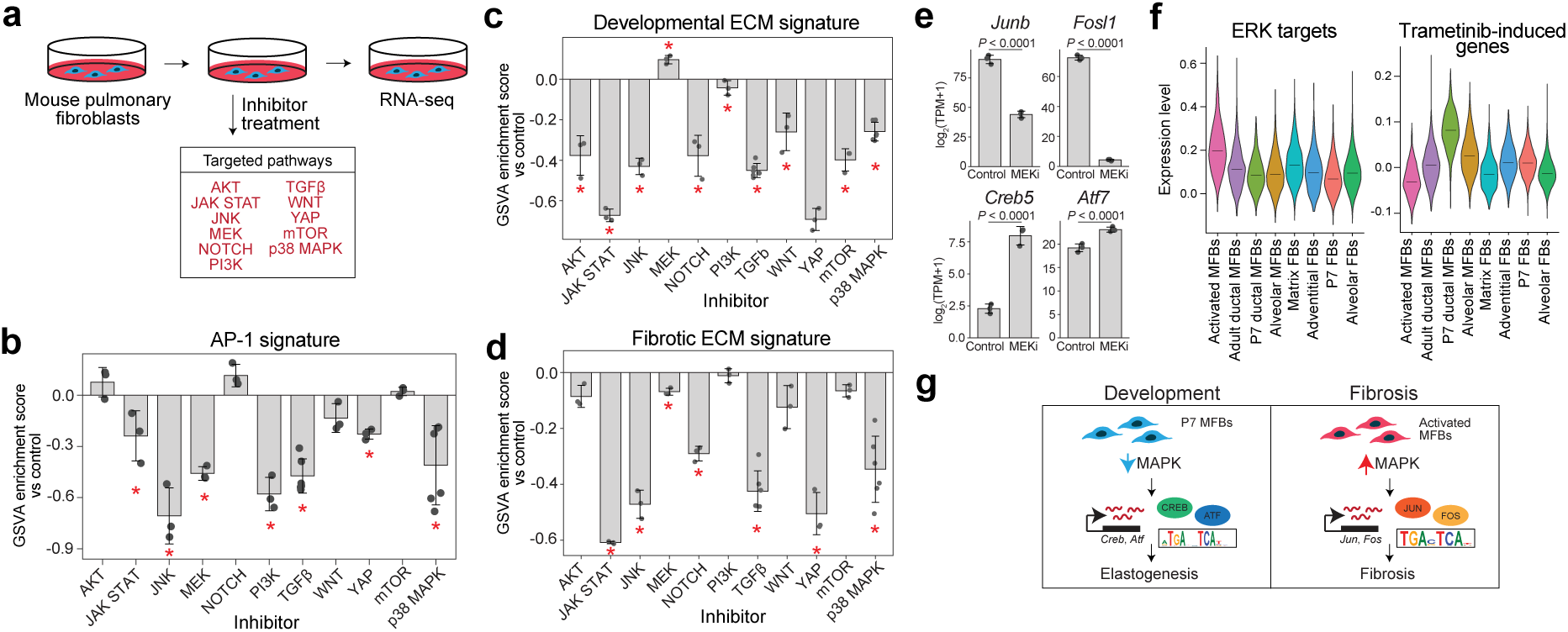
MAPK/ERK signaling regulates AP-1 activity. (a) Schematic of small molecule inhibitor experiment. MPFs were treated with inhibitors targeting the listed pathways for 24 hours before RNA-seq (*N* = 3 per treatment). (b-d) Bar graphs of GSVA enrichment scores of (b) an AP-1 target gene signature, (c) the developmental ECM signature, and (d) the fibrotic ECM signature in MPFs treated with listed inhibitors relative to the matched DMSO control (Supplementary Table 1). (e) Bar graphs of expression of TRE and CRE binding AP-1 family members in samples treated with the MEK inhibitor (MEKi) Trametinib and the matched DMSO control. (f) Violin plots of expression of (left) an ERK target gene signature and (right) the top upregulated genes with Trametinib treatment across fibroblast populations in the multiome dataset (Supplementary Table 1). (g) Model of AP-1 activity in P7 and activated myofibroblasts. Bar graphs show mean ± standard deviation. Violin plots show the median and the relative frequency of cells at a given expression value. *q* values were calculated by ordinary one-way ANOVA with two-stage linear step-up correction (Benjamini, Krieger and Yekutieli) (b-d, within-experiment), and *P* values were calculated by DESeq2 Wald test (e). * indicates *q* < 0.05.

### Pathological fibroblasts in human ILD upregulate AP-1 TRE activity

While we identified AP-1-mediated TRE activity as a driver of fibrotic ECM in mouse lung injury, its clinical relevance in humans remained to be established. To address this, we interrogated expression of the pathological TRE-driven ECM gene signature across single-cell RNA-seq datasets of human ILD^27,59,60^. Compared to control fibroblasts, the fibrotic ECM signature expression was elevated in fibroblasts across multiple types of ILD - including IPF, chronic hypersensitivity pneumonitis (cHP), and non-specific interstitial pneumonia (NSIP) - but not COPD, a disease characterized by airway and alveolar destruction rather than fibrosis (Fig. 7a-d, Supp. Fig. 7a-b). Meanwhile, the development-associated ECM signature was not significantly induced in IPF fibroblasts (Supp. Fig. 7c-f). In IPF, myofibroblasts drove expression of the fibrotic ECM transcriptional signature (Fig. 7e-g, Supp. Fig. 7g). In particular, pathological *CTHRC1+* fibroblasts displayed elevated program expression (Fig. 7f-g)^27,33,34^.

**Figure 7:**
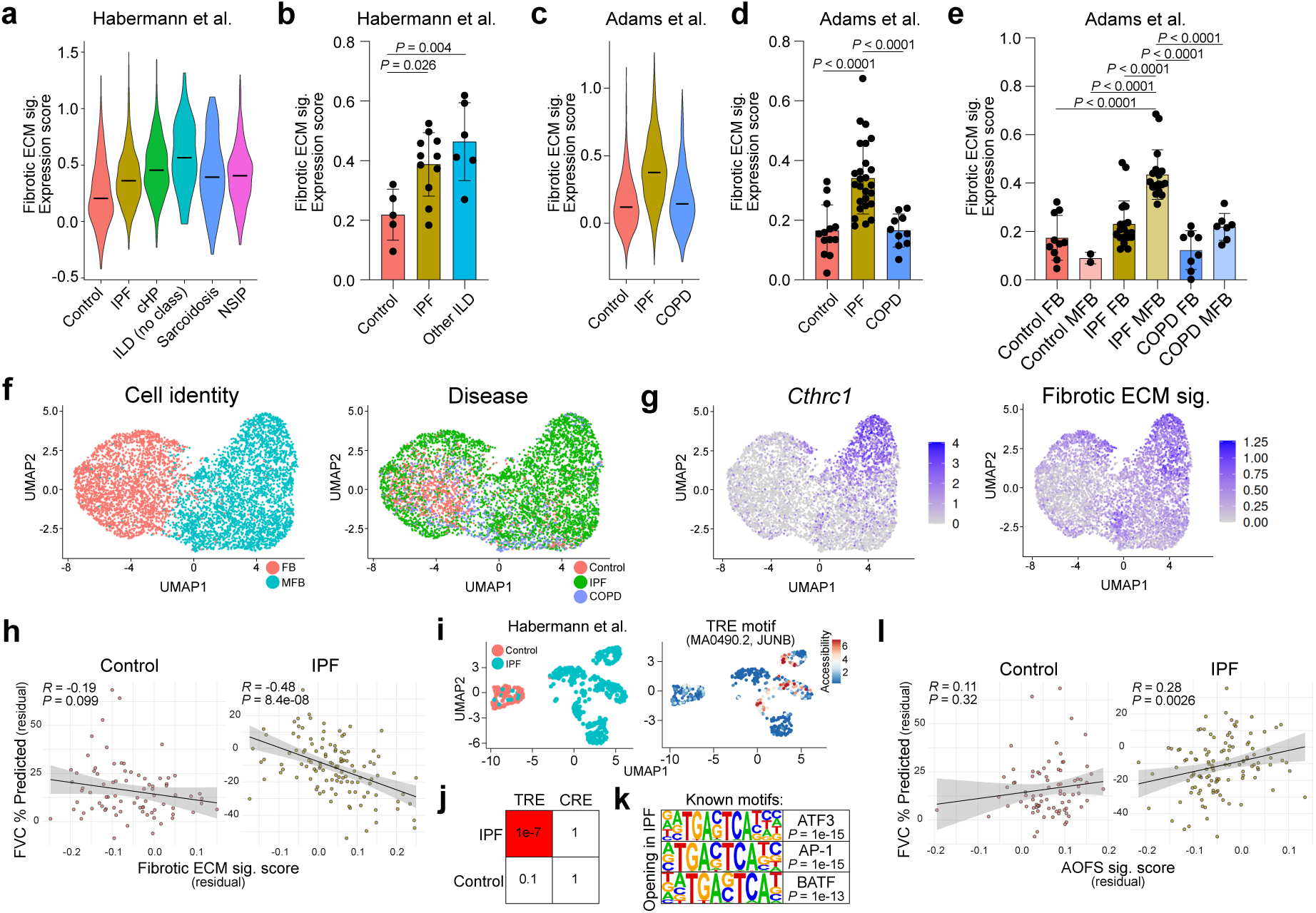
Fibrotic AP-1 activity is upregulated in human ILD. (a) Violin plot of expression of the fibrotic ECM program in all fibroblasts from a scRNA-seq dataset of control and ILD patients^59^. IPF, idiopathic pulmonary fibrosis; cHP, chronic hypersensitivity pneumonitis; ILD, interstitial lung disease; NSIP, non-specific interstitial pneumonia. (b) Bar graph of (a) pseudobulked by patient. Other ILD: cHP, ILD (no class), sarcoidosis, and NSIP samples. (c) Violin plot of expression of the fibrotic ECM program in all fibroblasts from a scRNA-seq dataset of control, IPF, and COPD patients^60^. (d) Bar graph of (c) pseudobulked by patient. (e) Bar graph of (d) split by expression in FBs and MFBs. (f) UMAP projections of all fibroblasts from (c) colored by (left) fibroblast type or (right) disease. (g) UMAP projections of expression of (left) an activated myofibroblast marker *CTHRC1* and (right) the fibrotic ECM signature. (h) Scatter plots of the adjusted fibrotic ECM program ssGSEA score versus FVC% predicted in (left) control and (right) IPF patients from a bulk transcriptomic dataset^61^. Rho (*R*), partial Spearman correlation coefficient. FVC, forced vital capacity. (i) UMAP plots of fibroblasts (alveolar FBs and MFBs) in a snATAC-seq dataset of control and IPF patients^62^ colored by (left) disease status or (right) motif accessibility (deviation score) of the JUNB motif. (j) *P*-value enrichment of TRE and CRE motifs in differentially accessible peaks open in control or IPF patient fibroblasts. (k) The top 3 known bZIP family member motifs by *P*-value enriched in differentially accessible peaks open in IPF patient fibroblasts identified by HOMER. (l) Scatter plots of the adjusted AFOS ECM program ssGSEA score versus FVC% predicted in (left) control and (right) IPF patients from a bulk transcriptomic dataset^61^. Violin plots show the median and the relative frequency of cells at a given expression value. Bar graphs show mean ± standard deviation. Scatter plots show a linear regression line with 95% confidence interval. *P* values were calculated by ordinary one-way ANOVA with Tukey’s multiple comparisons test (b, d, e), by Spearman’s rank correlation test (h, l), and by cumulative hypergeometric test (j, k, HOMER).

To test if fibrotic ECM program expression correlated with clinical disease severity, we queried a large-scale clinical cohort comprising microarray gene expression profiles from over 500 whole lung samples from control, ILD, and COPD patients with matched lung function metrics^61^. Consistent with the single-cell data, the fibrotic ECM program was upregulated in IPF and not COPD patients compared to control patients (Supp. Fig. 7h-i). To determine whether this signature expression was associated with lung function, multivariate regression analysis adjusting for age, sex, smoking status, and estimated fibroblast abundance using a pan-fibroblast gene expression score was performed. Increased expression of the fibrotic ECM gene expression program significantly correlated with reduced forced vital capacity (FVC) and reduced diffusing capacity for carbon monoxide (DLCO) in IPF patients (Fig. 7h, Supp. Fig. 7j-m). This suggests that elevated expression of this TRE-driven program was associated with declining lung function in IPF.

To further investigate AP-1 activity in IPF, TRE and CRE motif accessibility was analyzed in alveolar fibroblasts and myofibroblasts from a single-nuclei ATAC-seq dataset of healthy donor and IPF lungs^62^. TRE motif accessibility was enriched in chromatin regions that gained accessibility in IPF fibroblasts compared to healthy controls (Fig. 7i-k, Supplementary Table 7). Meanwhile, the CRE motif was not enriched in differential ATAC peaks in healthy nor diseased fibroblasts (Fig. 7j). We also inferred JUN/FOS transcription factor activity from the expression of their target regulons, which generally trended higher in IPF fibroblasts relative to controls (Supp. Fig. 7n)^63,64^. Together, these results implicate elevated AP-1 TRE motif activity in pathological fibroblasts.

AFOS expression inhibited AP-1 TRE activity and induced favorable transcriptional and cellular phenotypes in cultured fibroblasts. To explore how inhibition of AP-1 activity may affect outcomes in human IPF, we generated an expression program of the top genes induced by AFOS expression and evaluated whether this signature was associated with readouts of lung function in the large-scale clinical cohort previously described^61^. This AFOS program, which represents suppression of AP-1 TRE activity, was suppressed in IPF compared to control patients (Supp. Fig. 7o-p). After accounting for clinical and fibroblast covariates, increased AFOS signature expression was correlated with improved FVC and DLCO specifically in IPF patients, suggesting that lower AP-1 activity may benefit patient lung function (Fig. 7l, Supp. Fig. 7q-t).

## Discussion

Previous studies have identified lung fibroblast populations with specialized functions, yet the regulatory logic driving distinct ECM transcriptional profiles remained poorly defined^27–31,35^. Here, we generated a multimodal single-nuclei RNA- and ATAC-sequencing atlas across distinct physiological and pathological contexts to identify transcriptional and epigenetic drivers of divergent ECM programs. We discovered that unique myofibroblast populations arise in development and disease, each expressing specialized ECM transcriptional programs. Further, we found that differential AP-1 transcription factor complex activity underlies distinct ECM-producing myofibroblast populations. Specifically, transcriptional activity at TRE and CRE elements delineate fibrotic and developmental myofibroblasts, respectively. Antagonism of the AP-1 TRE signaling axis using a dominant-negative FOS protein was sufficient to repress the fibrotic ECM program and induce a more developmental ECM state. MEK inhibition produced ECM transcriptional phenotypes similar to AFOS expression and repressed transcription of TRE-binding AP-1 family members. These results reveal AP-1 transcription factor complex as a master orchestrator of myofibroblast ECM programs, with human ILD datasets confirming its relevance to disease and nominating it as a therapeutic target in lung fibrosis. Beyond the AP-1 axis described here, this atlas spans all major lung cell types across four biological contexts, enabling analogous studies in the epithelial, immune, and endothelial compartments.

### Distinct myofibroblasts in disease and development drive ECM synthesis

Across unique biological and ECM contexts, we identified fibroblast populations with distinct ECM roles. At homeostasis, adult ductal myofibroblasts, which align with elastic fibers at alveolar entrances, exhibited minimal expression of ECM transcriptional programs. Matrix fibroblasts, also present in the adult lung, displayed modestly elevated levels of ECM genes, such as *Col4a1* and *Fn1*, relative to alveolar fibroblasts. However, these cells lacked expression of myofibroblast markers, suggesting they mediate basal ECM turnover required for physiological maintenance of the healthy, uninjured lung. Notably, this matrix fibroblast population expanded with age and may therefore mediate the slight increase in fibrosis observed in normal aging^8,10^.

Pathological, activated *Cthrc1*+ *Spp1*+ myofibroblasts emerged within two weeks of bleomycin treatment and markedly upregulated fibrosis-associated transcriptional programs relative to other fibroblast populations. We identified an ECM gene expression program specifically induced in these activated myofibroblasts. Upregulation of this signature was consistent across multiple models of mouse lung injury and was recapitulated in human ILDs, including IPF. This established the fibrotic ECM signature as a conserved, core myofibroblast program downstream of diverse fibrotic stimuli. The absence of signature induction in COPD further marks this program as a signature of active fibrogenesis rather than a universal feature of lung disease or injury. Consistent with this observation, genes in the fibrotic ECM program were associated primarily with collagen biosynthesis, crosslinking, and ECM remodeling.

Myofibroblasts in the neonatal lung were distinct from those in the adult and diseased lung. Two myofibroblast populations have been previously described during alveolarization, referred to as alveolar/partitioning and ductal/entrance myofibroblasts^4,29,36–41,65^. Alveolar myofibroblasts are positioned at the leading edges of nascent secondary septa, where they deposit elastic ECM to form the structural scaffold for alveolar morphogenesis^4,29,38–41,65^. Ductal myofibroblasts are localized to alveolar entrances and synthesize the thick ring of elastic fibers that support ductal openings^29,65^. We discovered that these two developmental myofibroblast populations expressed a shared ECM program centered on elastogenesis, including genes essential for elastic fiber assembly and atypical collagens. This physiological program was not associated with injury or fibrosis, establishing it as a distinct developmental ECM state.

Thus, activated and developmental myofibroblasts express mutually exclusive, context-specific ECM programs with divergent functional consequences. These results suggest that activated myofibroblasts in disease do not re-engage developmental programs and instead produce tissue-inappropriate, fibrotic ECM. Because developmental myofibroblasts are uniquely specialized for elastic fiber synthesis, shifting myofibroblast identity from a pathological towards a developmental state may both inhibit fibrosis and restore physiological ECM. To identify mechanisms underlying these myofibroblast and ECM programs, we investigated the transcription factors driving these states.

### Differential AP-1 activity underlies myofibroblast identity

Our multiomics approach allowed us to predict differential transcription factor activity across cell populations. Strikingly, motif discovery analysis identified AP-1 activity in both pathological and developmental myofibroblasts. Closer examination revealed a single base pair difference between the AP-1-associated motifs of the two populations: activated myofibroblasts had increased accessibility at TRE motifs (5′ - <u>TGA</u>S<u>TCA</u> - 3′) whereas developmental myofibroblasts showed elevated accessibility at CRE motifs (5′ - <u>TGA</u>CG<u>TCA</u> - 3′). Marker genes and ECM-program genes of activated and developmental myofibroblasts were enriched for TRE and CRE motifs, respectively.

Integrating motif enrichment, expression profiling, and GRN analyses suggested that this differential motif preference was driven by context-dependent AP-1 family member activity. TRE activity in activated myofibroblasts after bleomycin treatment was likely driven by JUN/FOS family members, specifically JUNB. *Junb* expression was elevated in activated myofibroblasts relative to developmental myofibroblasts and the predicted JUNB GRN was likewise most highly expressed in this population. Interestingly, *Junb* was upregulated in cultured lung fibroblasts after AP-1 inhibition via AFOS expression, revealing a potential compensatory mechanism to restore AP-1 signaling and further supporting JUNB as a central mediator of TRE activity in activated myofibroblasts.

Meanwhile, the AP-1 family members CREB5, ATF2, and ATF7 emerged as candidate drivers of CRE motif activity in developmental myofibroblasts, showing increased expression and motif accessibility in developmental compared to activated myofibroblasts. CREB5 in particular was identified in our GRN analysis as driving CRE motif accessibility and transcriptional activity in P7 myofibroblasts. Interestingly, CREB5 was predicted to directly regulate *Eln*, a hallmark of developmental myofibroblast identity^3^. In cultured fibroblasts, CREB5 overexpression combined with *Junb* knockdown was sufficient to induce elastic fiber-associated genes, including *Eln*, in fibroblasts. These results, along with the specificity of *Creb5* expression in developmental myofibroblasts, nominate CREB5 as a candidate driver of elastic ECM deposition in the lung.

The differential activity of the AP-1 complex at TRE and CRE motifs between myofibroblast populations suggests that AP-1 serves as a molecular switch between distinct ECM transcriptional programs based on dimer composition and motif affinity. Our finding that dominant-negative inhibition of the TRE axis was sufficient to both repress the fibrotic ECM program and induce the developmental one demonstrated that AP-1 activity regulates these central biosynthetic programs. Interestingly, increased CRE motif accessibility was not necessary for induction of the developmental ECM program. Regulatory regions of these ECM genes may already have been available for binding, requiring no change in accessibility. Indeed, we found CRE motifs within invariant peaks associated with genes in the developmental signature, including *Mfap2*, *Col27a1*, and *Dst*. These results suggest that these regulatory regions were already accessible in cultured myofibroblasts, but transcription required a shift in the pool of AP-1 family members available for dimerization. The capacity of the AP-1 transcription factor complex to coordinate such divergent ECM programs establishes it as a master orchestrator of myofibroblast ECM states. Upstream of AP-1, the MAPK/ERK pathway emerged as a candidate regulator of this switch: MEK inhibition repressed expression of fibrotic programs and several TRE binding AP-1 family members while concurrently upregulating elastic ECM programs, positioning MAPK/ERK as a control point for AP-1 motif preference.

### Evolutionary and clinical implications of AP-1 mediated ECM regulation

Our findings suggest that lung fibroblasts exploit the inherent modularity of the AP-1 complex to shift between distinct transcriptional identities. The combinatorial diversity of the AP-1 family allows cells to generate a wide array of transcriptional outputs from a relatively small set of transcription factors and share regulatory networks^17,19,20^. The CRE-mediated program that underlies developmental myofibroblasts is associated with elastogenesis while the TRE-mediated program represents a collagen-centric wound healing response. Such flexibility may reflect an evolutionary history of subfunctionalization, allowing a single regulatory hub to be tuned across tissue contexts rather than requiring dedicated pathways for each ECM outcome. Within a tissue, multiple ECM programs could provide survival advantages. The complex ECM of the lung is meticulously constructed throughout development to meet the organ’s functional requirements, whereas rapid restoration of structural integrity via a dense collagenous network is preferable when repairing a wound. However, in fibrotic disease, the continued synthesis of tissue-inappropriate ECM becomes maladaptive, compromising lung function.

Across the adult, aged, neonatal, and injured lung, TRE and CRE motif opening was observed exclusively in activated and developmental myofibroblasts with one exception. TRE accessibility was also elevated in *Cldn4+* pre-alveolar type I transitional cells, which arise in response to alveolar injury and can persist in a non-regenerative state in disease^43,47,48^. These cells have been shown to exhibit pro-fibrotic signaling, displaying elevated expression of ECM genes such as *Col1a1* and *Itga2* compared to alveolar type 2 epithelial cells^43,47,66,67^. Consistent with our findings, TRE motif accessibility in these transitional cells was recently reported by another group, who suggest that JUNB and FOSB drive TRE motif opening and that attenuating *Junb*, *Fos*, and *Fosb* expression reduces fibroblast-activating senescence signaling from these cells^66^. The similarity in TRE-driven signaling across epithelial and mesenchymal cell types suggests a conserved response to injury in the lung that converges on ECM deposition and fibrosis.

Our analyses of TRE activity in human lung disease further underscored program conservation across species and the clinical relevance of this AP-1 regulatory axis. The fibrotic ECM signature was specifically enriched in myofibroblasts across multiple ILD subtypes, including IPF and cHP. TRE activity was concurrently elevated in IPF fibroblasts, suggesting that the fibrotic AP-1 signaling module is similarly active in pathological fibroblasts from both mouse and human lung disease. Consistent with this, recent work identified elevated TRE motif accessibility in fibroblasts from systemic sclerosis-associated ILD, extending the TRE signaling paradigm across ILDs^68^. Our AFOS results demonstrate that inhibiting TRE activity is sufficient to repress the pathological, fibrotic ECM signature, induce the physiological, developmental ECM program, and suppress type I collagen precursor production. Therapeutic interventions aimed at shifting

## Methods

### Animal studies

All mice were maintained in a barrier facility with a 12:12 hour light:dark cycle and allowed ad libitum access to food and water. Adult and aged C57BL/6J mice (Jackson Laboratory, catalog #000664) were shipped from Jackson Laboratory and acclimated for at least 7 days before experimentation. P7 neonatal mice were generated by breeding C57BL/6J mice in the barrier facility. All animal experiments were approved by the Institutional Animal Care and Use Committee at Calico Life Sciences. Four mice (two male and two female) were used per condition (adult untreated, adult saline-treated, adult bleomycin-treated, aged untreated, and P7 neonatal) in the single-nuclei multiomics experiment. Adult, aged, and neonatal lungs were harvested by perfusing with ice cold 1x PBS before flash freezing in liquid nitrogen.

### Bleomycin treatment

16 week old mice were anesthetized using isoflurane at 2.5-3.5%, and 50 ul of saline or bleomycin (1 U/kg, Meitheal Pharmaceuticals, catalog #092125) was delivered via intratracheal instillation. Animals were kept in a heated cage until ambulatory before transfer to their home cage and monitored for 2 weeks after treatment. At 13 days post-treatment, µCT imaging was performed. At 14 days after treatment, animals were euthanized, and lungs were perfused with ice cold 1x PBS, dissected, and flash frozen in liquid nitrogen.

### micro-CT imaging

Lung structure was assessed using micro-CT (μCT) imaging. Mice were anesthetized with 1-3% isoflurane with medical air. Breathing rates were maintained between 50-60 breaths per minute by adjusting the isoflurane concentration. Focused lung scans were acquired with a MILabs micro-CT system (U-CT, MILabs, Utrecht, Netherlands) using the following settings: mouse ultra focus gated protocol, X-ray voltage 50 kV, current 130 µA, exposure 75 ms, binning of 2X2, 0.6 degrees step angle, 4 projections per angle. Radiation delivered per scan was less than 1 Gy with an imaging time of approximately 7 minutes. Micro-CT images were reconstructed using vendor provided software at 50 µm isotropic resolution with respiratory gating. Micro-CT imaging was conducted 13 days post saline or bleomycin delivery as well as prior to tissue collection for adult and aged mice.

Automatic lung segmentation was performed using a U-Net model trained on manual lung segmentations. An internal reference atlas with individual lobe segmentations was registered to segmented lungs using Elastix1 (Version 5.2, rigid and affine registration). Masks for lung lobes (Left, right cranial, right middle, right caudal and right accessory) were propagated to the original image space and used for lobe specific analysis. Image intensity thresholds were used to define fibrosis or inflammation (> −250 HU), normal lung (−750 to −250) and emphysema (< −750 HU). Various lung parameters like end-expiratory lung volume, high signal intensity volume (“fibrosis volume”), low signal intensity volume (emphysema volume), percentage of lung affected by fibrosis or emphysema were computed for each lobe.

### Single-nuclei RNA- and ATAC-sequencing and analysis

Flash frozen lungs samples from adult, aged, neonatal, saline-treated, and bleomycin-treated mice were used for the single-nuclei RNA- and ATAC-sequencing multiomics experiment. Nuclei were isolated from frozen tissues using the Chromium Nuclei Isolation with RNase Inhibitor Kit (10x Genomics, catalog #1000494) according to the manufacturer’s protocol. Nuclei were counted using a NucleCounter NC-3000 (Chemometec), and 15,400 nuclei were used to generate the single cell gene expression and ATAC libraries using the 10x Genomics Chromium Next GEM Single Cell Multiome ATAC + Gene Expression Kit (10x Genomics, catalog #1000285, #1000234, #1000215). Libraries were sequenced on a NovaSeq6000. The resulting data was processed through bcl2fastq and further processed through cellranger-arc count (10x Genomics). The resulting gene expression matrices and ATAC counts were loaded into R (v4.3.2) and processed using Seurat (v4.3.0.1, https://github.com/satijalab/seurat) and Signac (v1.11.0, https://github.com/timoast/signac) for downstream analyses^69,70^. Cells were filtered to only include those with the following parameters: (1) more than 200 but fewer than 25,000 RNA counts, (2) more than 200 but fewer than 2,500 RNA features, (3) less than 10% mitochondrial reads, (4) more than 700 but fewer than 100,000 ATAC counts, (5) nucleosome signal less than 2, and (6) transcription start site enrichment greater than 2.

Gene expression matrices were processed using SCTransform() to normalize, scale, find variable features, and regress the variable number of RNA counts and percent mitochondrial reads. Principal component analysis was run to identify the first 50 major axes of variation using RunPCA(). DNA accessibility data were processed by finding the most frequently observed features using FindTopFeatures(), computing the term-frequency inverse-document frequency using RunTFIDF(), and running singular value decomposition using RunSVD(). Batch integration was then performed using Harmony on both the gene expression and ATAC datasets (v1.2.3, https://portals.broadinstitute.org/harmony)^71^. The variable used in group.by.vars was the sampleID of each individual sample. Joint UMAP visualization of gene expression and chromatin accessibility data was performed via the weighted nearest neighbor (WNN) method in Seurat using FindMultiModalNeighbors(). In brief, the WNN graph was constructed using the Harmony-integrated pca and lsi reductions of the gene expression and chromatin accessibility data. Clustering was performed with the WNN graph using FindClusters() and visualized by UMAP dimensionality reduction via RunUMAP(). Cell identity was determined through analyses of gene and ATAC peak markers of each cluster using FindAllMarkers() and by mapping previously defined expression and accessibility patterns of cells in the lung. To perform PC analysis of gene expression profiles, pseudobulk profiles were generated for each sample by averaging the SCT-normalized expression data across all cells belonging to that sample. The 2,000 most variable genes across pseudobulk profiles were selected and used for PCA. Based on these results, untreated and saline-treated 16 week old adult mice were combined together. Cell cycle scoring was performed using the Seurat implementation of the cell cycle scoring function, as previously described^72^. Fibroblasts were subsetted, and the subsequent gene expression and DNA accessibility matrices were processed again as described above, with the addition of cell cycle scoring was also included as a variable to regress in SCTransform(). Fibroblast identity was determined through analyses of gene and ATAC peak markers of each cluster using FindAllMarkers() and by mapping previously defined expression and accessibility patterns of cells in the lung. Motif accessibility scores were calculated using ChromVAR (v1.22.1, https://github.com/GreenleafLab/chromVAR)^46^. Gene module scores of ECM programs were calculated using AddModuleScore() on the normalized and scaled RNA assay. Pseudobulk profiles of fibroblast populations were generated by grouping cells by sample and specific fibroblast type, retaining only groups with at least 30 cells. Raw RNA counts were aggregated using AggregateExpression() and gene signature module scores were summarized by averaging per-cell scores within each group.

The fibrotic and developmental ECM signatures were developed by identifying differentially expressed genes between P7 myofibroblasts (alveolar and ductal) and activated myofibroblasts using FindMarkers(). The developmental ECM signature was created by taking significantly upregulated genes in developmental myofibroblasts identified from this analysis that overlapped with terms found in the Reactome “Extracellular Matrix Organization” gene set (R-HSA-1474244). The fibrotic ECM signature was created by taking significantly upregulated genes in activated myofibroblasts from this analysis that overlapped with terms found in the Reactome “Extracellular Matrix Organization” gene set (R-HSA-1474244). To increase specificity of the fibrotic ECM signature, this initial gene set was then filtered to only include genes that were identified as upregulated in activated myofibroblasts compared to the rest of the fibroblast population using FindMarkers().

Differentially expressed peaks were identified across fibroblast populations using FindAllMarkers(). Transcription factor motif analyses were performed individually on differentially accessible peaks that increased in accessibility in individual cell populations using the HOMER motif discovery tool (v4.11.0, https://homer.ucsd.edu/homer) with the findMotifsGenome() command^44^. All identified peaks across fibroblasts were used as background peaks.

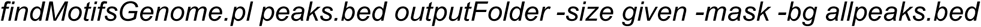

ATAC peaks associated with genes in the ECM programs were identified by identifying differentially expressed peaks between P7 myofibroblasts and activated myofibroblasts using FindMarkers(), taking peaks with significantly increased accessibility in each population, and using GREAT (Genomic Regions Enrichment of Annotations Tool, v4.0.4, https://great.stanford.edu) to link peaks to genes^73,74^. The resulting peaks associated with ECM program genes were used in HOMER analyses.

Enhancer-mediated gene regulatory networks (GRNs) were inferred from the multiome data using GRaNIE (v1.4.1, https://github.com/bioconductor-source/GRaNIE)^49^. RNA and ATAC counts were pseudobulked across fibroblast cell types using AggregateExpression(). Pseudobulked matrices were added to the GRN object with quantile normalization (limma) for RNA and DESeq2 size-factor normalization for ATAC. Transcription factor binding sites were defined from HOCOMOVO v12 (H12INVIVO) PWMScan motifs and overlapped with accessible peaks. Transcription factor-peak links were computed by Pearson correlation between transcription factor expression and peak accessibility, and peak-gene links by Pearson correlation within a 250 kilobase window around each gene. The network was filtered at an FDR threshold of 0.3 for both connections (Benjamini-Hochberg). Inferred target genes of AP-1 family transcription factors were extracted from the filtered GRN and used to construct gene expression signatures. Gene module scores of these programs were calculated using AddModuleScore(). Transcription factor-centric regulatory networks were visualized as directed graphs linking each transcription factor to its top correlated target genes (FDR < 0.25). GRN-derived gene signature module scores were pseudobulked by sample and cell type as described above. The regulatory connection between CREB5 and *Eln* was visualized by plotting the CREB5 GRN-linked peaks and gene body at the *Eln* locus.

### Analysis of published mouse single-cell RNA-sequencing datasets

The gene expression matrices from Curras-Alonso et al. (GSE211713), Tsukui et al. (GSE132771), Zepp et al. (GSE149563), Strunz et al. (GSE141259), and Narvaez del Pilar et al. (GSE180822) were loaded into R (v4.3.2) and processed using Seurat (v4.3.0.1, https://github.com/satijalab/seurat) for downstream analyses^27,29,32,35,43^. Gene expression matrices were processed using SCTransform() to normalize, scale, find variable features, and regress the variable number of RNA counts. Principal component analysis was run to identify the first 50 major axes of variation using RunPCA(). Batch integration was performed using Harmony (v1.2.3, https://portals.broadinstitute.org/harmony). The variable used in group.by.vars was the sample ID of each individual mouse. Clustering was performed by constructing a nearest neighbor graph using FindNeighbors() and identifying clusters of cells by a shared nearest neighbor modularity optimization-based clustering algorithm using FindClusters(). Clusters were visualized by UMAP dimensionality reduction via RunUMAP(). *Col1a1*+ clusters were subsetted, and gene expression matrices were processed again as described above. Gene module scores of ECM programs were calculated using AddModuleScore() on the normalized and scaled RNA assay. For the Curras-Alonso et al. (GSE211713), Tsukui et al. (GSE132771), and Strunz et al. (GSE141259) datasets, pseudobulk profiles were generated by grouping cells by sample, retaining only groups with at least 30 cells. Gene signature module scores were summarized by averaging per-cell scores within each group. For the Curras-Alonso et al. (GSE211713) and Tsukui et al. (GSE132771) datasets, differentially expressed genes between untreated and bleomycin-treated or 4 month irradiation-treated *Col1a1*+ subsetted cells were identified using FindMarkers(). The Curras-Alonso fibrosis ECM signature was identified by taking significantly upregulated genes in the 4 month irradiation condition identified from this analysis that overlapped with terms found in the Reactome “Extracellular Matrix Organization” gene set (R-HSA-1474244). The Tsukui fibrosis ECM signature was identified by taking significantly upregulated genes in the bleomycin condition identified from this analysis that overlapped with terms found in the Reactome “Extracellular Matrix Organization” gene set (R-HSA-1474244).

### Cell culture

Adult mouse pulmonary fibroblast cells (ScienCell, catalog #M3300-57) were maintained in Fibroblast Medium (FM, Sciencell, catalog #2301) supplemented with fetal bovine serum (Sciencell, catalog #0010), fibroblast growth supplement (Sciencell, catalog #2352), and Antibiotic solution (penicillin streptomycin, Sciencell, catalog #0503) in an incubator set to 37°C, 20% O_2_, and 5% CO_2_. Cells were routinely tested for mycoplasma and always tested negative. Cells were grown on plastic (polystyrene) plates of various sizes (Falcon). When cells reached 70-80% confluence, cells were passaged by washing the plate with PBS and then incubating with TrypLE Express Enzyme (Gibco, catalog #12604) until cells lifted from the plate. Media was added to neutralize the reaction and the cell suspension was spun at 300 x *g* for 5 minutes. The resulting cell pellet was resuspended in media and counted using a ViCell XR Cell Analyzer with Trypan Blue staining to assess cell number and viability (Beckman Coulter). Cells were used in experiments at passage 2. For cell counting experiments, the ViCell XR Cell Analyzer with Trypan Blue was used to quantify cell number. Pro-Col1a1 was quantified using the Mouse Pro-Collagen I alpha 1 ELISA kit (Abcam, catalog #ab210579). Cells were counted and spun at 300g for 5 minutes at 4C, then washed 3x with 1x PBS. After the last wash, supernatant was removed and cells were flash frozen in LN2. Samples were thawed on wet ice for 1 minute before performing sample extraction and the assay procedure as described. For drug treatments, cells were seeded 24 hours prior to treatment. Cells were then treated with compounds for 24 hours before harvest. Treatments were done across three rounds of experiments with a separate DMSO control for each experiment. All compounds were purchased from TargetMol. Experiment 1: CIL56 (1 μm, catalog #T4309), Galunisertib (10 μm, catalog #T2510), MK-2206 (3 μm, catalog #T1952), SB-431542 (10 μm, catalog #T1726), STX-0119 (25 μm, catalog #T60160), Tanzisertib (10 μm, catalog #T14895), Wnt-C59 (1 μm, catalog #T2242). Experiment 2: Alpelisib (5 μm, catalog #1921), Doramapimod (10 μm, catalog #T6277), Rapamycin (5 μm, catalog #T1537), Skepinone-L (1 μm, catalog #T6130). Experiment 3: Trametinib (50 nm, catalog #T2125), RO4929097 (10 μm, catalog #T6274).

### Electroporation

siRNA and expression constructs were delivered to mouse pulmonary fibroblasts via electroporation using the 4D-Nucleofector System with the X Unit (Lonza). The P1 Primary Cell 4D-Nucleofector X Kit S (Lonza, catalog #V4XP-1032) and the P1 Primary Cell 4D-Nucleofector X Kit L (Lonza, catalog #V4XP-1012) were used for electroporation. Cells were grown to 80% confluence. Plates were washed with PBS and then incubated with TrypLE Express Enzyme (Gibco) until cells lifted from the plate. Media was added to neutralize the reaction. Cells were counted twice using a ViCell XR Cell Analyzer with Trypan Blue staining to assess cell viability (Beckman Coulter). For reactions in the Nucleocuvette Strips (Kit S), 1×10^5^ cells with 0.4 ug and/or 30 nM siRNA were used. For reactions in the Single Nucleocuvettes (Kit L), 1×10^6^ cells with 0.2 ug of plasmid and/or 30 nM siRNA were used. EGFP (pRP[Exp]-EF1A>EGFP), AFOS (pRP[Exp]-CMV>{A-FOS}), Creb5 (pRP[Exp]-EF1A>mCreb5[NM_001327821.1]), and Atf7 (pRP[Exp]-EF1A>mAtf7[NM_001310066.1]) overexpression constructs were purchased from VectorBuilder. The sequence for AFOS was obtained from the Vinson Lab CMV500 A-FOS plasmid^75^. ON-TARGETplus siRNA SMARTpools targeting mouse Jun and Fos family members and the ON-TARGETplus non-targeting pool were purchased from Horizon Discovery. Cells were spun at 90 x *g* for 10 minutes and resuspended in the 4D-Nucleofector P1 solution, siRNA and/or plasmid substrates were added, and mastermixes were transferred to the Nucleocuvette vessels. Vessels were placed in the 4D-Nucleofector Core Unit and run on the DS-137 protocol. Vessels were incubated at room temperatures for 10 minutes before plating.

### Bulk RNA-sequencing and analysis

For bulk RNA-sequencing experiments, cells were lysed directly on cell culture plates in Buffer RLT Plus (Qiagen, catalog #1053393). The lysate was homogenized using Qiashredder columns (Qiagen, catalog #79656) and stored at −80°C. Once all samples were collected, RNA was isolated using an RNeasy mini kit (Qiagen, catalog #74134). Quality and concentration of RNA were determined using a 2100 Bioanalyzer Instrument (Agilent). cDNA libraries were constructed using the NEBNext Ultra II Directional RNA Library Prep Kit for Illumina with the polyA mRNA workflow (New England Biolabs, catalog #E7760). Samples were sequenced using a NovaSeq6000 (Illumina). Raw sequencing reads were demultiplexed with bcl2fastq. Reads were trimmed with cutadapt (v2.8.0,, https://github.com/marcelm/cutadapt) with parameters -j 8 -e 0.1 -q 20 -O 1 and aligned to build GRCm39 of the mouse genome using STAR (v2.7.6a, https://github.com/alexdobin/STAR) with extra parameters --twopassMode Basic --outSAMunmapped Within --outSAMattributes NH HI AS NM MD --outSAMstrandField intronMotif^76,77^. Alignments were quantified using all features in the Gencode version M34 GTF (https://www.gencodegenes.org/mouse/release_M34.html) with featureCounts from subread (v1.6.2, https://subread.sourceforge.net) in paired end mode with options -O –fraction. Differentially expressed genes (DEGs) were identified using DESeq2 (v1.50.2, https://github.com/thelovelab/DESeq2) in R (v4.5.3) with a significance cutoff of *P*-adjusted < 0.05^78^. For the RNA-seq experiment with AFOS, AFOS was compared to the EGFP sample. The DESeq2 Wald test was used to calculate fold change and *P* values using the model: ∼condition, where condition is a variable capturing the overexpression construct used. For the RNA-seq experiment manipulating individual AP-1 family members, samples with knockdown of individual AP-1 family members were compared to the siControl sample, while samples with overexpression or combined knockdown/overexpression of AP-1 family members were compared to the EGFP sample. The DESeq2 Wald test was used to calculate fold change and *P* values using the model: ∼condition, where condition is a variable capturing the knockdown or overexpression performed. For the small molecule inhibitor experiments, treatments were performed across several rounds of experiments and sequencing runs, each with a DMSO-matched control. Each experiment was done as a separate sequencing run. Read counts from the experiments were merged by intersecting gene sets and analyzed together in DESeq2. The DESeq2 Wald test was used to calculate gene-level fold changes and *P* values using the model ∼experiment + condition, where condition captures the small molecule inhibitor perturbation and experiment is a batch term capturing the experiment of origin, with the pooled DMSO conditions as the reference.

Principal component analysis was performed using DESeq2. Heatmap visualization was performed using pheatmap (v1.0.12, https://github.com/raivokolde/pheatmap). Enrichr (v3.2.0, https://maayanlab.cloud/Enrichr) was used to perform Reactome gene ontology analyses^79–81^. Composite gene expression signature scores were calculated for each sample by multiplying the centered log_2_ fold change of each gene in the signature in each sample by the DESeq2 stat value for each gene. Cell type deconvolution of bulk RNA-seq data was performed using MuSiC (v1.0.0, https://github.com/xuranw/MuSiC) and the single-nuclei gene expression data generated in this study as the reference^82^. Analysis was restricted to genes detected in both the bulk and single-nuclei datasets. Cell type proportions were estimated using weighted non-negative least squares via the music_prop() function. These values were then averaged across all genes in the signature in each sample to generate the signature score. ssGSEA scores were calculated using the GSVA package (v2.4.8, https://github.com/rcastelo/GSVA) with the ssgseaParam function applied to variance-stabilized transformed counts from DESeq2^83^. Cell type gene signatures were defined using the top 10 marker genes for each fibroblast subtype identified by FindAllMarkers() in Seurat in the single-nuclei dataset generated in this study. GSVA enrichment scores were calculated using the gsvaParam function from the same package, applied to variance-stabilized transformed counts. Pathway gene sets were obtained from MSigDB via msigdbr (v26.1.0, https://github.com/igordot/msigdbr) or are listed in Supplementary Table 1. For display, scores were centered on the matched control — each experiment’s own vehicle control in the small molecule screen, and siControl or EGFP as appropriate in the knockdown/overexpression experiment — by subtracting the mean control score for the corresponding signature, so that bar heights represent the mean change relative to control. For drug treatment GSVA analyses, because treatments were done across three experiments each with its own control, scores were analyzed separately within each experiment: for each signature, an ordinary one-way ANOVA was fit across the conditions of a given experiment, and each inhibitor condition was compared to the control from that same experiment. Multiple comparisons were corrected by controlling the false discovery rate using the two-stage linear step-up procedure of Benjamini, Krieger and Yekutieli (Q = 5%), applied within each experiment; *q* values are reported. Statistical testing was performed on the unnormalized enrichment scores. To generate the Trametinib-induced gene signature and plots of AP-1 family member expression with MEK inhibition, the Trametinib-treated samples and matched DMSO controls were analyzed separately in DESeq2 using the model ∼condition, with the DESeq2 Wald test used to calculate fold changes and *P* values. The top 200 significantly upregulated genes by fold change in the Trametinib condition were used for the signature.

To generate the correlation scatter plots comparing AFOS overexpression and MEK inhibition, the small molecule inhibitor and AFOS RNA-seq experiments were merged by intersecting gene sets and read counts were analyzed together with DESeq2. Counts were combined into a single DESeq2 object for normalization and variance stabilization. The control conditions from all experiments were treated as a common reference group. Counts were variance-stabilized and experiment-of-origin was modeled as a batch term and removed from the variance-stabilized expression matrix with limma::removeBatchEffect() (v3.66.0, https://bioconductor.org/packages/release/bioc/html/limma.html), supplying condition as the design matrix so that differences between conditions were preserved^84^. The batch-corrected values were then averaged across the replicates of each condition to give a single expression profile per condition. For each gene, log_2_ fold changes relative to the common control were calculated by subtracting the control profile from the MEK inhibition and AFOS condition profiles, respectively. Gene-wise concordance between the MEK inhibition and AFOS responses was assessed by plotting these log_2_ fold changes against one another and computing Spearman’s rank correlation using cor.test() in R. This was performed across three gene sets: the 500 most variable genes (by variance across condition-averaged profiles), the mouse orthologues of the Reactome “Extracellular Matrix Organization” gene set (REACTOME_EXTRACELLULAR_MATRIX_ORGANIZATION), obtained from MSigDB (C2:CP:REACTOME) via msigdbr (v26.1.0, https://github.com/igordot/msigdbr), and the combined genes of the developmental and fibrotic ECM signatures.

### Bulk ATAC-sequencing and analysis

For the bulk ATAC-sequencing experiment, cells were harvested from plates using TrypLE Express Enzyme as described above and frozen in liquid nitrogen in FM with 10% DMSO at a density of 1×10^6^ cells/mL. Samples were sent to Admera Health for ATAC library prep and sequencing. Samples were sequenced using a NovaSeq6000 (Illumina). FASTQ files were trimmed and low quality reads were filtered using cutadapt (v2.8.0, https://github.com/marcelm/cutadapt) to remove primer sequences.

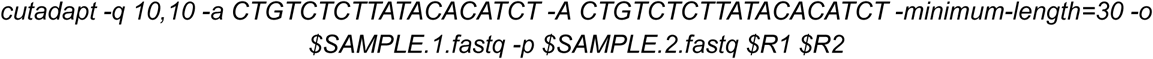

Trimmed FASTQ files were then aligned to the genome using bowtie2 with the extra parameter-X 1000 (v2.3.4, https://github.com/BenLangmead/bowtie2), and sorted based on genomic location and indexed using Samtools (v1.14, https://github.com/samtools/samtools)^85–88^. Quality control, peak calling, peak-gene associations, differential peak analysis, and principal component analyses were performed using ChrAccR (v0.9.21, https://github.com/GreenleafLab/ChrAccR) in R (v4.3.2). Default parameters were used to generate the promoter peaks list. To generate the genome-wide peaks list (all peaks) via MACS2 (v2.2.9.1, https://pypi.org/project/MACS2), the setConfigElement doPeakCalling() was used with annotationPeakGroupAgreePerc() = 0.5. Transcription factor motif analyses were performed individually on differentially accessible peaks with increased accessibility from each comparison using the HOMER motif discovery tool (v4.11.0, https://homer.ucsd.edu/homer) with the findMotifsGenome() command as described above. All identified peaks across conditions were used as background peaks. TRE and CRE motif enrichment were calculated using custom motif files in the -mknown argument of the findMotifsGenome() command.

| >TGAGSTCA TRE 7.0 |  |  |  |
| --- | --- | --- | --- |
| 0.0 | 0.0 | 0.0 | 1.0 |
| 0.0 | 0.0 | 1.0 | 0.0 |
| 1.0 | 0.0 | 0.0 | 0.0 |
| 0.0 | 0.5 | 0.5 | 0.0 |
| 0.0 | 0.0 | 0.0 | 1.0 |
| 0.0 | 1.0 | 0.0 | 0.0 |
| 1.0 | 0.0 | 0.0 | 0.0 |

| >TGACGTCA CRE 8.0 |  |  |  |
| --- | --- | --- | --- |
| 0.0 | 0.0 | 0.0 | 1.0 |
| 0.0 | 0.0 | 1.0 | 0.0 |
| 1.0 | 0.0 | 0.0 | 0.0 |
| 0.05 | 0.9 | 0.0 | 0.05 |
| 0.05 | 0.0 | 0.9 | 0.05 |
| 0.0 | 0.0 | 0.0 | 1.0 |
| 0.0 | 1.0 | 0.0 | 0.0 |
| 1.0 | 0.0 | 0.0 | 0.0 |

TRE and CRE peak annotation and distance from peak center were calculated with HOMER using annotatePeaks(). Enrichr (v3.2.0, https://maayanlab.cloud/Enrichr) was used to perform Reactome gene ontology analyses using genes linked to peaks by ChrAccR.

### Analysis of published human single-cell RNA-sequencing datasets

The RDS object from Habermann et al. (GSE135893) was loaded into R (v4.3.2) and processed using Seurat (v4.3.0.1, https://github.com/satijalab/seurat) for downstream analyses^59^. Fibroblast clusters were subset using previously defined cell type labels. Gene module scores of ECM programs were calculated using AddModuleScore() on the normalized and scaled RNA assay. Pseudobulk profiles were generated by grouping cells by sample or sample and cell type, retaining only groups with at least 20 cells. Gene signature module scores were summarized by averaging per-cell scores within each group. Samples were either grouped by control samples and specific disease types or control, IPF, and broad ILD samples (comprising cHP, ILD [no class], sarcoidosis, and NSIP). Control and IPF samples were further subsetted. Transcription factor activity was inferred per cell by applying VIPER (v1.34.0, https://bioconductor.org/packages/viper) via decoupleR (v2.6.0, https://github.com/saezlab/decoupleR) to the scaled RNA expression data using DoRothEA (v1.12.0, https://github.com/saezlab/dorothea) regulons (confidence levels A–C)^63,64,89^.

The gene expression matrices from Adams et al. (GSE136831) were loaded into R (v4.3.2) and processed using Seurat (v4.3.0.1, https://github.com/satijalab/seurat) for downstream analyses^60^. Fibroblast clusters were subset using previously defined cell type labels. Gene expression matrices were processed using SCTransform() to normalize, scale, find variable features, and regress the variable number of RNA counts. Principal component analysis was run to identify the first 50 major axes of variation using RunPCA(). Batch integration was performed using Harmony (v1.2.3, https://portals.broadinstitute.org/harmony). The variable used in group.by.vars was the sample ID of each individual mouse. Clustering was performed by constructing a nearest neighbor graph using FindNeighbors() and identifying clusters of cells by a shared nearest neighbor modularity optimization-based clustering algorithm using FindClusters(). Clusters were visualized by UMAP dimensionality reduction via RunUMAP(). Gene module scores of ECM programs were calculated using AddModuleScore() on the normalized and scaled RNA assay. Pseudobulk profiles were generated by grouping cells by sample or sample and cell type, retaining only groups with at least 20 cells. Gene signature module scores were summarized by averaging per-cell scores within each group.

The gene expression matrices from Tsukui et al. (GSE132771) were loaded into R (v4.3.2) and processed using Seurat (v4.3.0.1, https://github.com/satijalab/seurat) for downstream analyses^27^. Gene expression matrices were processed using SCTransform() to normalize, scale, find variable features, and regress the variable number of RNA counts. Principal component analysis was run to identify the first 50 major axes of variation using RunPCA(). Batch integration was performed using Harmony (v1.2.3, https://portals.broadinstitute.org/harmony). The variable used in group.by.vars was the sample ID of each individual mouse. Clustering was performed by constructing a nearest neighbor graph using FindNeighbors() and identifying clusters of cells by a shared nearest neighbor modularity optimization-based clustering algorithm using FindClusters(). Clusters were visualized by UMAP dimensionality reduction via RunUMAP(). *COL1A1*+ clusters were subsetted, and gene expression matrices were processed again as described above. Gene module scores of ECM programs were calculated using AddModuleScore() on the normalized and scaled RNA assay. Pseudobulk profiles were generated by grouping cells by sample or sample and cell type, retaining only groups with at least 20 cells. Gene signature module scores were summarized by averaging per-cell scores within each group.

### Analysis of published human single-nuclei RNA and ATAC-sequencing datasets

The gene expression matrices and fragments files from Valenzi et al. (GSE214085) were loaded into R (v4.3.2) and processed using Seurat (v4.3.0.1, https://github.com/satijalab/seurat) and Signac (v1.11.0, https://github.com/timoast/signac) for downstream analyses^62^. The snRNA-seq dataset was processed by normalizing, finding variable features, and scaling the RNA assay. Principal component analysis was run to identify the first 50 major axes of variation using RunPCA(). Batch integration was performed using Harmony (v1.2.3, https://portals.broadinstitute.org/harmony). The variable used in group.by.vars was the sample ID of each individual mouse. Clustering was performed by constructing a nearest neighbor graph using FindNeighbors() and identifying clusters of cells by a shared nearest neighbor modularity optimization-based clustering algorithm using FindClusters(). Clusters were visualized by UMAP dimensionality reduction via RunUMAP(). Fibroblast clusters were identified through analysis of fibroblast marker expression. DNA accessibility data were processed by finding the most frequently observed features using FindTopFeatures(), computing the term-frequency inverse-document frequency using RunTFIDF(), and running singular value decomposition using RunSVD(). Batch integration was performed using Harmony (v1.2.3, https://portals.broadinstitute.org/harmony). Clustering was performed as described above. A gene activity matrix was computed by summing chromatin accessibility over gene bodies and promoter regions using GeneActivity(). The fibroblast cell-type label was transferred from the annotated scRNA-seq reference to the snATAC-seq cells using FindTransferAnchors() and TransferData(). The predicted fibroblasts in the ATAC data were subset and the accessibility data was processed as described above. Motif accessibility scores were calculated using ChromVAR (v1.22.1, https://github.com/GreenleafLab/chromVAR). Differentially expressed peaks between control and IPF fibroblasts were identified using FindMarkers(). Transcription factor motif analyses were performed individually on differentially accessible peaks that increased in accessibility in individual cell populations using the HOMER motif discovery tool (v4.11.0, https://homer.ucsd.edu/homer) with the findMotifsGenome() command. Differentially expressed peaks were identified across fibroblast populations using FindAllMarkers(). Transcription factor motif analyses were performed on differentially accessible peaks that increased in accessibility in IPF samples using the HOMER motif discovery tool (v4.11.0, https://homer.ucsd.edu/homer) with the findMotifsGenome() command as described above. All identified peaks across fibroblasts were used as background peaks. TRE and CRE motif enrichment were calculated using custom motif files in the -mknown argument of the findMotifsGenome() command as described above.

### Analysis of published human microarray data of chronic lung disease

Gene expression data from the Lung Genomics Research Consortium (GSE47460) were downloaded via GEOquery (v2.78.0, https://github.com/seandavi/GEOquery) and analyzed in R (v4.5.3)^61,90^. The GPL14550 (Agilent 8x60K whole-genome microarray) was used, comprising 213 control and patient samples. Samples were classified into four broad disease groups: control, IPF, COPD, and other ILD (including NSIP, HP, RB-ILD, DIP, ILD associated with CVD, and unclassifiable ILD). Probes were mapped to gene symbols using the platform feature data, and multi-gene probe entries were resolved by retaining the first listed symbol. Probes mapping to the same gene were collapsed by averaging using limma::avereps() (v3.66.0, https://bioconductor.org/packages/release/bioc/html/limma.html).

ssGSEA was performed on the full gene expression matrix using the GSVA package (v2.4.8, https://github.com/rcastelo/GSVA) to score each sample for the fibrotic ECM program, the AFOS gene signature, and a fibroblast abundance signature. The AFOS gene signature was created using the top 10 genes upregulated in the AFOS condition in the EGFP versus AFOS comparison by *P*-adjusted value in the bulk RNA-sequencing experiment. The fibroblast abundance signature was created by using fibroblast population markers identified in Figure 2a of Tsukui et al^27^. To assess the contribution of potential confounding variables to the expression of these signatures, a multivariable linear regression model was fit separately for each signature in control and IPF samples, using the signature ssGSEA score as the outcome and age, sex, smoking status, and fibroblast abundance score as predictors. Sex was coded as female versus male and smoking status as never- versus ever-smoker; age and fibroblast abundance score were standardized to unit standard deviation prior to model fitting to compare effect sizes across predictors on a common scale. Samples with missing covariate data were excluded. Regression coefficients with 95% confidence intervals were visualized as forest plots, and *P* values were obtained from two-sided *t-*tests of each coefficient without correction for multiple comparisons. To assess the association between signature expression and lung function independently of confounders, partial correlations between each signature and %predicted FVC or %predicted DLCO were computed in control and IPF samples by regressing each variable on age, sex, smoking status (never versus ever/current), and fibroblast abundance score using lm(), then computing Spearman’s rank correlation between the two sets of residuals. Covariates were included on their original scales for this analysis.

### Statistical analysis

Data analysis and statistical tests were performed using GraphPad Prism software (v.10.2.0) and in R (v4.3.2 and v4.5.3). All statistical tests are denoted in the figure legends.

## Data and code availability

Data from the single-nuclei RNA- and ATAC-sequencing multiomics experiment, bulk RNA-seq experiments, and the bulk ATAC-seq experiment are available at GEO under the accession number GSE346825. Previously published datasets used in this paper are available at GEO under the accession numbers: GSE135893, GSE136831, GSE132771, GSE214085, GSE149563, GSE141259, GSE211713, GSE180822, and GSE47460. Code for processing and analyzing data modalities and generating figures is available at https://github.com/calico/ap1-lung-paper/releases/tag/v0.9.0.

## Acknowledgements

We thank S. Nazertehrani and E. Karlsson for organizing and performing breedings for the P7 lungs; D. Hendrickson, J. Brown, and the Calico Genomics lab for their help in processing sequencing samples and guidance on data analysis; and A. Gillich and J. Arron for advice and critical reading of the manuscript.

## Contributions

A.M.K designed and performed experiments, interpreted and analyzed data, and wrote the manuscript. K.S., B.M-N., W.C., and J.P. performed the bleomycin experiment. K.S. and G.D. performed µCT imaging experiments. A.L. harvested samples for the single-nuclei multiomics experiment. A.C. and J.R. contributed to the design of the animal studies. J.G.R. designed and interpreted experiments and wrote the manuscript. All authors reviewed and approved the final manuscript for submission.

## Funding

This study was funded by Calico Life Sciences LLC.

## Competing interests

This research was funded by Calico Life Sciences LLC, South San Francisco, CA, where all authors were employees at the time the research was conducted. The authors declare no other competing financial conflicts.

## Supplemental files

**Supplemental Figures:** Seven Supplemental Figures (S1-S7) with legends as one PDF.

**Supplemental Table 1:** Gene expression signatures

**Supplemental Table 2:** HOMER results from Figure 4b

**Supplemental Table 3:** HOMER results from Figure 4c

**Supplemental Table 4:** HOMER results from Supplementary Figure 4e

\**Supplemental Table 5:** GRaNIE GRN network output

**Supplemental Table 6:** HOMER results from Fig. 5c

**Supplemental Table 7:** HOMER results from Figure 7k

## Notes

https://www.ncbi.nlm.nih.gov/geo/query/acc.cgi?acc=GSE346825

