## Supplementary Figures for "AP-1 specifies developmental versus fibrotic extracellular matrix transcriptional programs in the lung"

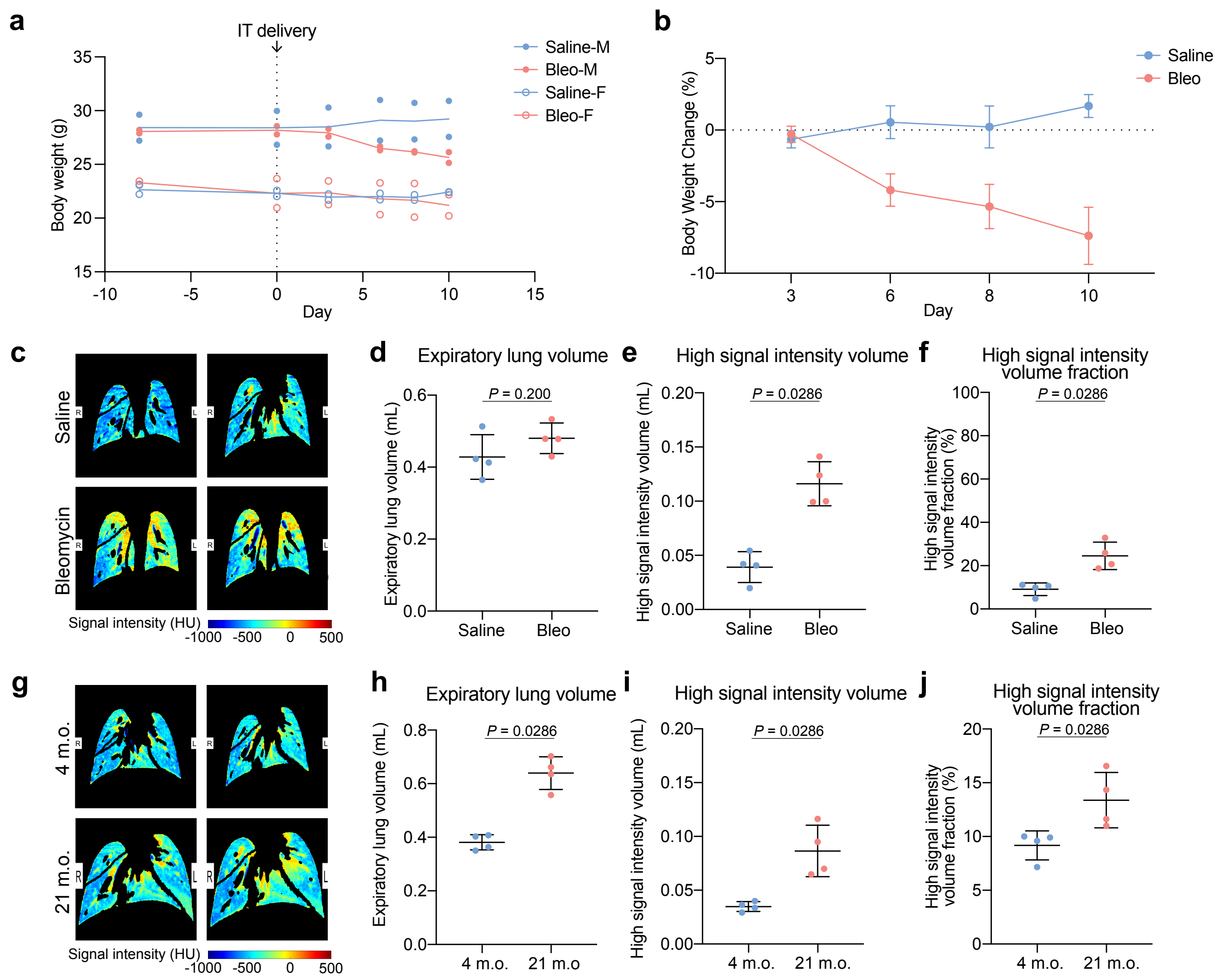


**Supplementary Figure 1: µCT and body weight confirm bleomycin-induced lung injury and age-associated changes.**

(a) Line graph of mouse body weight over time before and after IT delivery of saline or bleomycin split by treatment condition and sex (*N* = 2 per group). Day 0 indicates the day of IT delivery. IT, intratracheal; M, male; F, female. (b) Line graph of the relative change in body weight over time after IT delivery of saline or bleomycin, split by treatment condition (*N* = 4 per group). Weights are all relative to day 0. (c) Representative µCT images of lungs 13 days post saline or bleomycin treatment. HU, Hounsfield units. (d-f) Bar graphs of (d) expiratory lung volume, (e) high signal intensity volume, and (f) high signal intensity volume fraction in saline and bleomycin-treated mice. (g) Representative µCT images of lungs from untreated 4 m.o. and 21 m.o. mice. (h-j) Bar graphs of (h) expiratory lung volume, (i) high intensity lung volume, and (j) high signal intensity volume fraction in 4 m.o. and 21 m.o. mice.

Line graphs (a-b) show mean ± standard error of the mean. Bar graphs (d-f, h-j) show mean ± standard deviation. *P* values were calculated by Mann-Whitney U Test (d-f, h-j).

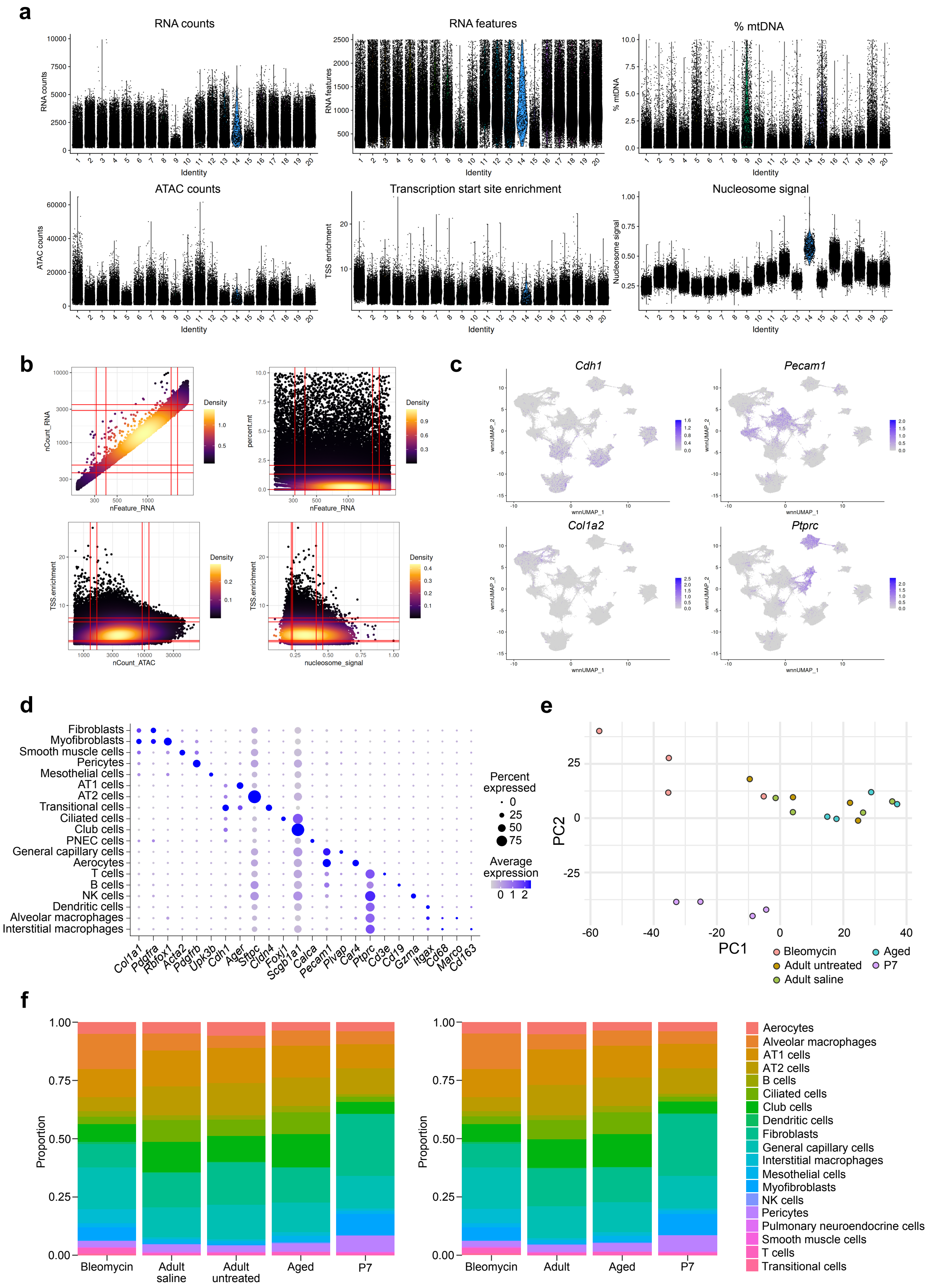

a
RNA counts
RNA features
% mtDNA
ATAC counts
Transcription start site enrichment
Nucleosome signal
Identity
TSS enrichment
b
nCount_RNA
nFeature_RNA
percent.mt
Density
TSS.enrichment
nCount_ATAC
nucleosome_signal
c
Cdh1
Pecam1
Col1a2
Ptprc
wnnUMAP_1
wnnUMAP_2
d
Fibroblasts
Myofibroblasts
Smooth muscle cells
Pericytes
Mesothelial cells
AT1 cells
AT2 cells
Transitional cells
Ciliated cells
Club cells
PNEC cells
General capillary cells
Aerocytes
T cells
B cells
NK cells
Dendritic cells
Alveolar macrophages
Interstitial macrophages
Col1a1 Pdgfra Rbfox1 Acta2 Pdgfrb Upk3b Cdh1 Ager Sftpc Cldn4 Foxj1 Scgb1a1 Calca Pecam1 Plvap Car4 Ptprc Cd3e Cd19 Gzma Itgax Cd68 Marco Cd163
Percent expressed
0
25
50
75
Average expression
0 1 2
e
PC1
PC2
Bleomycin
Adult untreated
Adult saline
Aged
P7
f
Proportion
Bleomycin
Adult saline
Adult untreated
Aged
P7
Adult
Aerocytes
Alveolar macrophages
AT1 cells
AT2 cells
B cells
Ciliated cells
Club cells
Dendritic cells
Fibroblasts
General capillary cells
Interstitial macrophages
Mesothelial cells
Myofibroblasts
NK cells
Pericytes
Pulmonary neuroendocrine cells
Smooth muscle cells
T cells
Transitional cells

**Supplementary Figure 2: snRNA- and ATAC-seq quality control and cell type definition.**
(a) Violin plots of RNA counts, RNA features, percent mitochondrial DNA, ATAC counts, transcription start site enrichment, and nucleosome signal in each individual sample after filtering. mt, mitochondrial. (b) Density plots of quality control variables in all cells. (c) UMAP projections of expression of cell type lineage markers *Cdh1*, *Pecam1*, *Col1a2*, and *Ptprc* across all cells after filtering. Lines indicate the 5th, 10th, 90th, and 95th quantiles. (d) Dot plot of expression of cell-type markers in all cells after filtering. (e) Principal component analysis (PCA) plot of samples colored after pseudobulking by gene expression. (f) Bar graphs displaying the relative proportion of cells in each cluster (left) before and (right) after merging the adult saline and adult untreated samples.

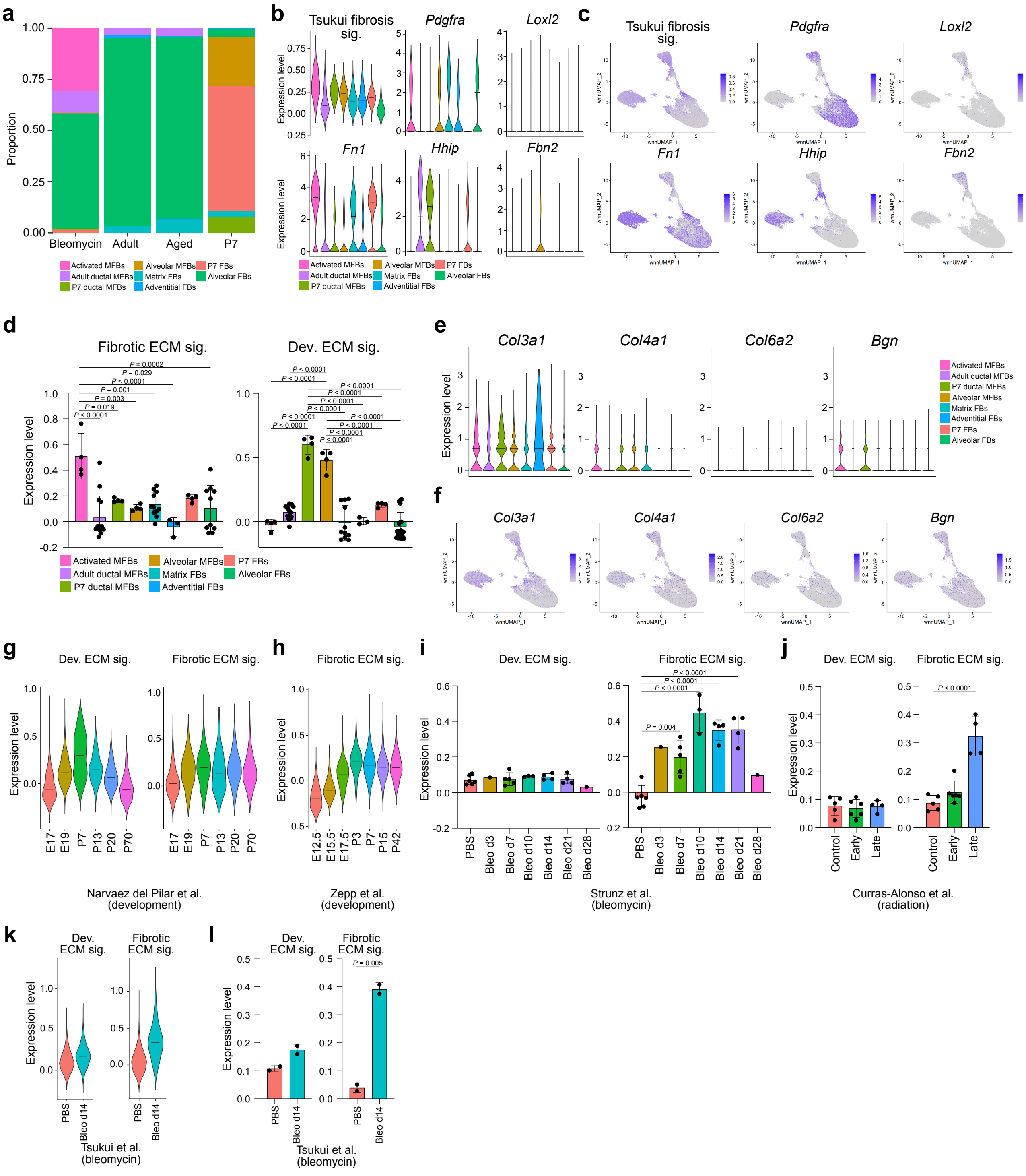


**Supplementary Figure 3: Fibroblast subtype characterization and cross-dataset validation of the fibrotic and developmental ECM programs.**

(a) Bar graphs displaying the relative proportion of each fibroblast cluster in each condition. (b) Violin plots of expression of a fibrosis gene signature derived from a bleomycin-induced lung fibrosis mouse model (Supplementary Table 1) and other individual gene markers of lung fibroblast subtypes across fibroblast populations[27]. (c) UMAP plot visualizations of (b). (d) Bar graphs of expression of the (left) fibrotic and (right) developmental ECM gene program in data from (Fig. 3a-b) pseudobulked by mouse. (e) Violin plots of expression of various ECM genes across fibroblast populations. (f) UMAP plot visualizations of (e). (g) Violin plot of expression of the (left) developmental and (right) fibrotic ECM programs in fibroblasts from a scRNA-seq dataset of mouse lung development[29]. Expression is shown in fibroblasts from E17.5 to P70. (h) Violin plot of expression of the fibrotic ECM program in fibroblasts from a scRNA-seq dataset of mouse lung development[35]. Expression is shown in fibroblasts from E12.5 to postnatal P42. (i) Bar graphs of the (left) developmental and (right) fibrotic ECM gene program in data from (Fig. 3d) pseudobulked by mouse[43]. (j) Bar graphs of the (left) developmental and (right) fibrotic ECM gene program in data from (Fig. 3e) pseudobulked by mouse[32]. (k) Violin plots of expression of the (left) developmental and (right) fibrotic ECM programs in a scRNA-seq dataset of bleomycin-induced mouse lung injury[27]. Expression is shown in fibroblasts from control (PBS-treated) lungs or from bleomycin-treated lungs (14 days post-treatment). (l) Bar graphs of the (left) developmental and (right) fibrotic ECM gene program in data from (k) pseudobulked by mouse.

Violin plots show the median and the relative frequency of cells at a given expression value. Bar graphs show mean ± standard deviation. *P* values were calculated by ordinary one-way ANOVA with Tukey's multiple comparisons test (d, i, j; for space reasons, only significant changes from activated myofibroblasts [d, left], P7 ductal and alveolar myofibroblasts [d, right], PBS [i], or control [j] are shown) and by two-tailed Welch's *t*-test (l).

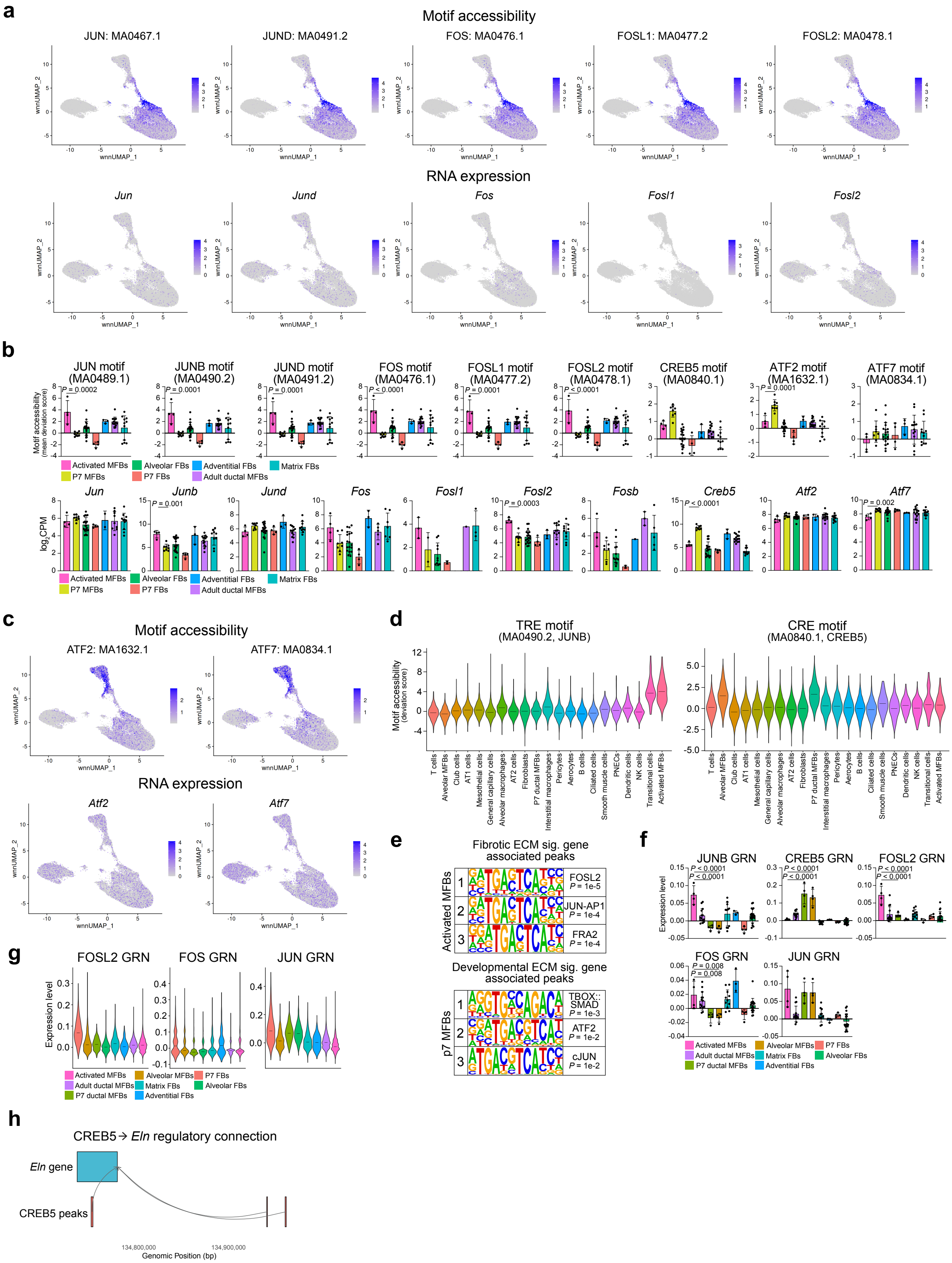

a
Motif accessibility
JUN: MA0467.1
JUND: MA0491.2
FOS: MA0476.1
FOSL1: MA0477.2
FOSL2: MA0478.1
wnnUMAP_1
wnnUMAP_2
RNA expression
Jun
Jund
Fos
Fosl1
Fosl2
b
JUN motif (MA0489.1)
JUNB motif (MA0490.2)
JUND motif (MA0491.2)
FOS motif (MA0476.1)
FOSL1 motif (MA0477.2)
FOSL2 motif (MA0478.1)
CREB5 motif (MA0840.1)
ATF2 motif (MA1632.1)
ATF7 motif (MA0834.1)
Motif accessibility (mean deviation score)
P = 0.0002
P = 0.0001
P < 0.0001
P = 0.002
Activated MFBs
P7 MFBs
Alveolar FBs
P7 FBs
Adventitial FBs
Adult ductal MFBs
Matrix FBs
Jun
Junb
Jund
Fos
Fosl1
Fosl2
Fosb
Creb5
Atf2
Atf7
log2CPM
P = 0.001
P = 0.0003
P < 0.0001
c
Motif accessibility
ATF2: MA1632.1
ATF7: MA0834.1
RNA expression
Atf2
Atf7
d
TRE motif (MA0490.2, JUNB)
CRE motif (MA0840.1, CREB5)
Motif accessibility (deviation score)
T cells
Alveolar MFBs
Club cells
AT1 cells
Mesothelial cells
General capillary cells
Alveolar macrophages
AT2 cells
Fibroblasts
P7 ductal MFBs
Interstitial macrophages
Pericytes
Aerocytes
B cells
Ciliated cells
Smooth muscle cells
PNECs
Dendritic cells
NK cells
Transitional cells
Activated MFBs
e
Fibrotic ECM sig. gene associated peaks
Activated MFBs
FOSL2 P = 1e-5
JUN-AP1 P = 1e-4
FRA2 P = 1e-4
Developmental ECM sig. gene associated peaks
p7 MFBs
TBOX::SMAD P = 1e-3
ATF2 P = 1e-2
cJUN P = 1e-2
f
JUNB GRN
CREB5 GRN
FOSL2 GRN
FOS GRN
JUN GRN
Expression level
P < 0.0001
P = 0.008
Activated MFBs
Adult ductal MFBs
P7 ductal MFBs
Alveolar MFBs
Matrix FBs
Adventitial FBs
P7 FBs
Alveolar FBs
g
FOSL2 GRN
FOS GRN
JUN GRN
Expression level
Activated MFBs
Adult ductal MFBs
P7 ductal MFBs
Alveolar MFBs
Matrix FBs
Adventitial FBs
P7 FBs
Alveolar FBs
h
CREB5→ Eln regulatory connection
Eln gene
CREB5 peaks
134,800,000
134,900,000
Genomic Position (bp)

**Supplementary Figure 4: AP-1 family member expression and activity varies across fibroblast populations.**

(a) UMAP plots visualizing (top) motif accessibility (deviation score) or (bottom) RNA expression ($log_2$ counts per million, CPM) of TRE binding AP-1 family members across fibroblast populations. (b) Bar graphs of (top) pseudobulked motif accessibility (deviation score) or (bottom) pseudobulked RNA expression of TRE and CRE binding AP-1 family members. Alveolar myofibroblast and P7 ductal myofibroblast samples are grouped into P7 myofibroblasts. (c) UMAP plots visualizing (top) motif accessibility (deviation score) or (bottom) RNA expression of CRE binding AP-1 family members across fibroblast populations. (d) Violin plots of (left) TRE (JUNB) and (right) CRE (CREB5) motif accessibility across all cell types. (e) The top 3 known motifs by *P*-value enriched in (top) peaks significantly increasing in accessibility in activated myofibroblasts linked to genes in the fibrotic ECM signature and in (bottom) peaks significantly increasing in accessibility in P7 myofibroblasts linked to genes in the developmental ECM signature by HOMER. (f) Bar graphs of pseudobulked AP-1 family member expression GRNs. (g) Violin plots of expression of the FOSL2, FOS, and JUN GRNs. (h) Cartoon visualizing the link between accessibility of the ATAC peaks on chromosome 5 to the *Eln* gene.

Violin plots show the median and the relative frequency of cells at a given expression value. Bar graphs show mean ± standard deviation. *P* values were calculated by ordinary one-way ANOVA with Tukey's multiple comparisons test (b, top, f; for space reasons, only significant changes between activated myofibroblasts and P7 myofibroblasts are shown), DESeq2 Wald test (b, bottom; for space reasons, only significant changes between activated myofibroblasts and P7 myofibroblasts are shown), and by cumulative hypergeometric test (e, HOMER).

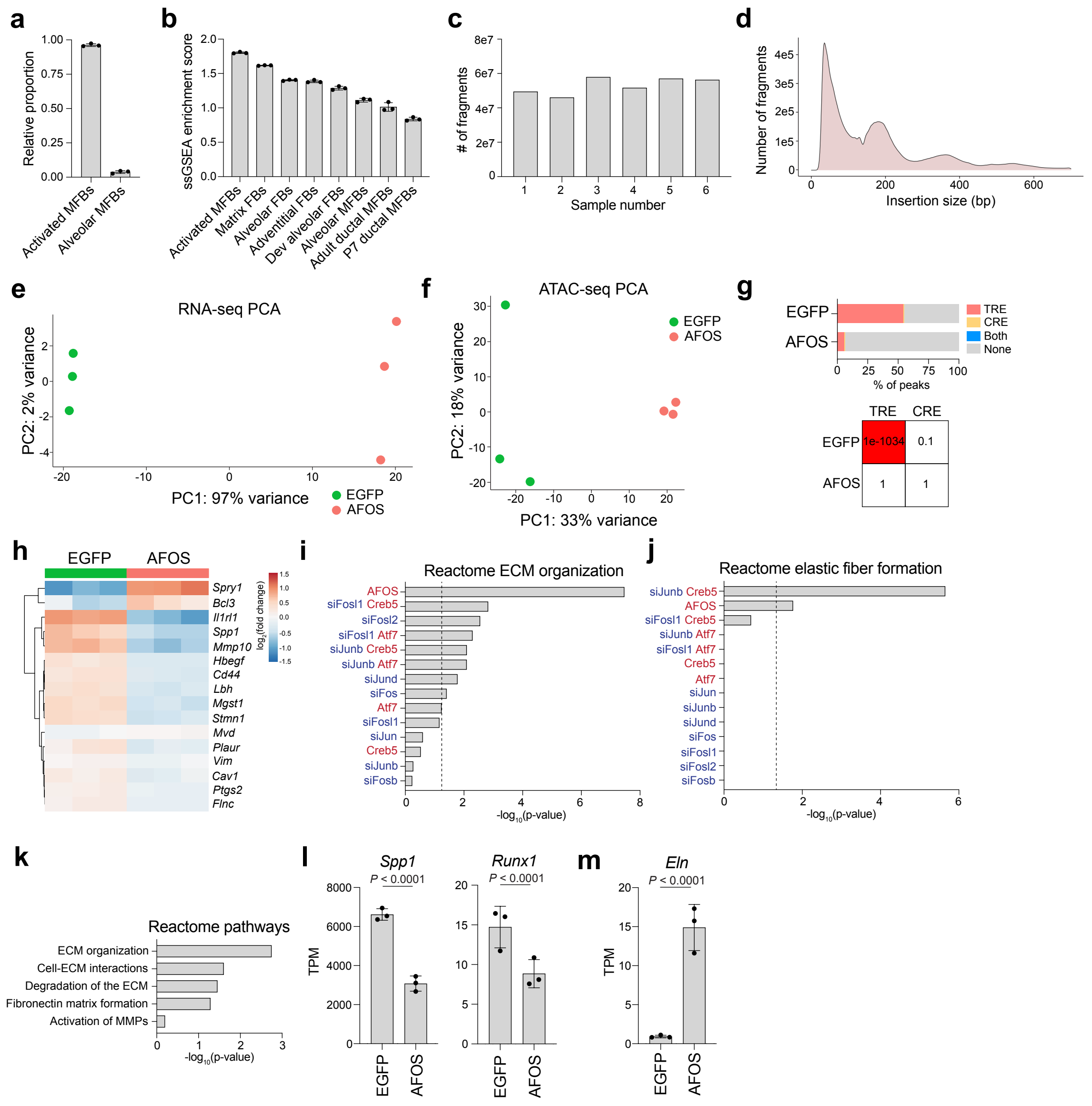

**Supplementary Figure 5: Supporting characterization of the MPF culture system and dominant-negative AFOS perturbation.**

(a) Bar graph of MuSiC-estimated cell-type proportions using bulk RNA-seq of cultured MPFs, deconvolved using the annotated multiome fibroblast populations as reference. (b) Bar graph of ssGSEA enrichment scores for the top 10 markers of each multiome fibroblast population, scored in bulk RNA-seq of cultured MPFs ($N$ = 3). (c) Bar graph of fragment count in each bulk ATAC-seq sample. (d) Representative histogram (sample 1) of fragment number versus insertion size for the ATAC-seq experiment. bp, base pair. (e-f) PCA plots of samples in the bulk (e) RNA- and (f) ATAC-seq experiments. (g) (Top) Bar graph of the proportion of differential peaks ($P$ < 0.05, peaks used in Figs. 5d-e) increasing in accessibility in each condition that contains TRE and CRE motifs and (bottom) the $P$-value enrichment of these motifs across conditions. (h) Heatmap of genes in the AP-1 target gene signature across samples. (i-j) Bar graph of Enrichr enrichment for the Reactome (i) "Extracellular Matrix Organization" or (j) “Elastic Fiber Formation” term among differentially expressed genes in each AP-1 perturbation versus its control (siControl for knockdowns; EGFP for overexpression and combination conditions). Dotted line marks *P* = 0.05. (k) Bar graph of enrichment of ECM-related Reactome gene ontology terms in genes linked with differentially accessible peaks (*P*-adjusted < 0.05). (l-m) Bar graphs of gene expression markers of (l) activated or (m) developmental myofibroblasts across conditions.

Bar graph (a, b, l, m) shows mean ± standard deviation. *P* values were calculated by cumulative hypergeometric test (g, HOMER) and by DESeq2 Wald test (l-m).

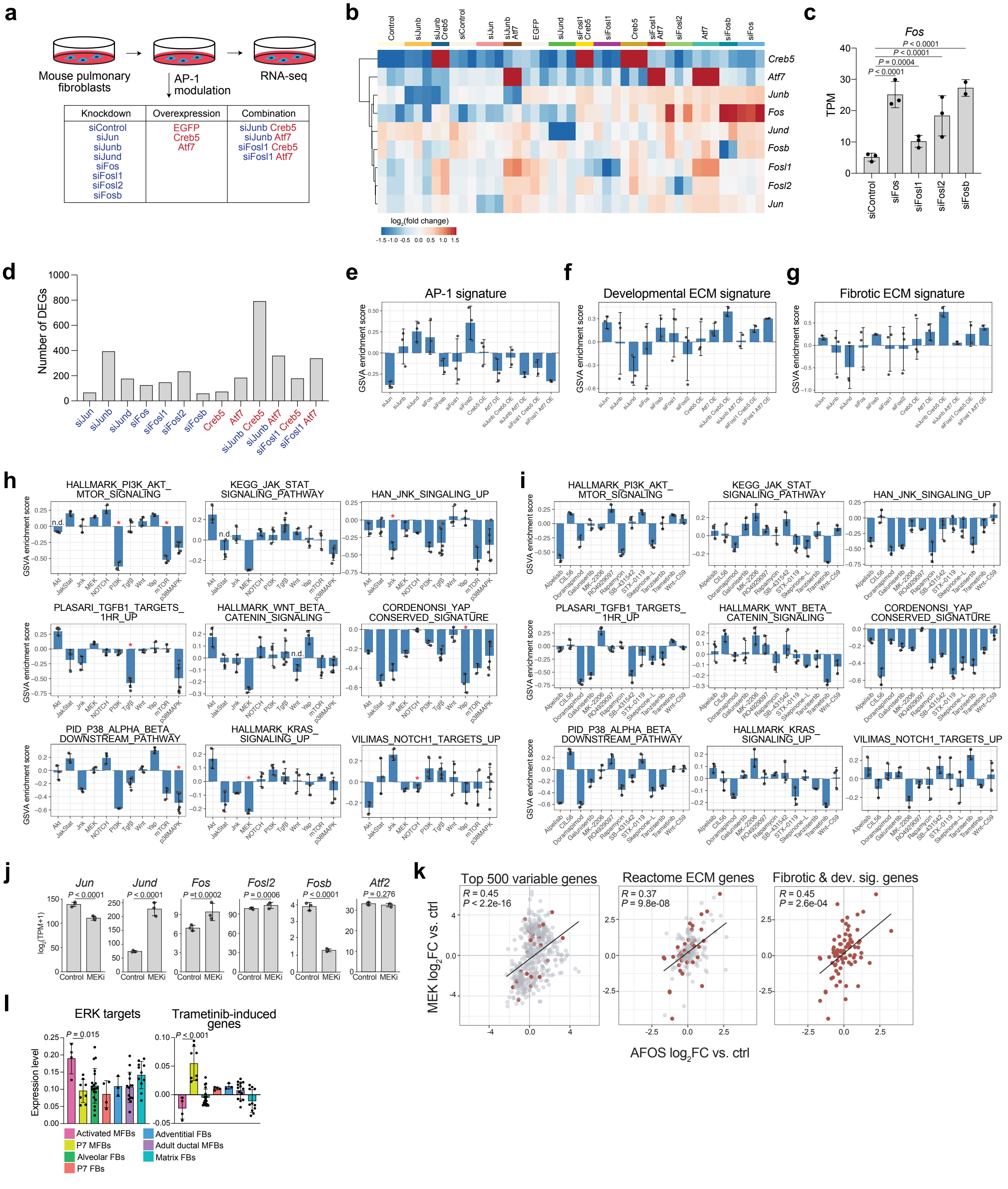


**Supplementary Figure 6: Modulating individual AP-1 family members and AP-1 regulating pathways in lung fibroblasts.**

(a) Schematic of AP-1 family member knockdown and overexpression experiment. siRNAs and expression constructs were delivered to MPFs by electroporation, and 24 hours after delivery samples were harvested for RNA-seq ($N$ = 3 per condition). (b) Heatmap of all AP-1 family members targeted for knockdown or overexpression in MPFs. (c) Bar graph of *Fos* expression in control samples and samples with knockdown of FOS family members. (d) Bar graph displaying the number of differentially expressed genes identified in the contrast of each AP-1 manipulation condition versus the matched control (siControl for individual knockdowns, EGFP for overexpression and combination samples). (e-g) Bar graphs of GSVA enrichment scores of (e) an AP-1 target gene signature, (f) the developmental ECM signature, or (g) the fibrotic ECM signature across inhibitor treatment conditions relative to the matched DMSO control (Supplementary Table 1). (h-i) Bar graph of GSVA enrichment scores of expression signatures of pathways targeted by the inhibitors from (Fig. 6a) across (h) broad inhibitor conditions or (i) individual drug treatments relative to the matched DMSO control. n.d., no discovery (j) Bar graphs of expression of TRE and CRE binding AP-1 family members in samples treated with Trametinib and the matched DMSO controls. (k) Gene-wise correlation of $\log_2$ fold changes (relative to the control) between MEK inhibition and AFOS expression across (left) the 500 most variable genes among all samples in the inhibitor and AFOS experiments, (middle) Reactome ECM organization genes, and (right) the developmental and fibrotic ECM signature genes. Genes from the developmental and fibrotic ECM signatures are highlighted in red. *R*, partial Spearman correlation coefficient. (l) Bar graphs of expression of (left) an ERK target gene signature and (right) the top upregulated genes with Trametinib treatment in data from (Fig. 6g) pseudobulked by mouse (Supplementary Table 1).

Bar graphs (c, e-j, l) show mean ± standard deviation. *P* values were calculated by DESeq2 Wald test (c, j), by Spearman's rank correlation test (k), and by ordinary one-way ANOVA with Tukey's multiple comparisons test (l; for space reasons, only significant differences between activated and P7 MFBs are shown). *q* values were calculated by ordinary one-way ANOVA with two-stage linear step-up correction (Benjamini, Krieger and Yekutieli) (h, within-experiment; only significant differences are shown for the condition(s) related to the gene signature). * indicates $q < 0.05$.

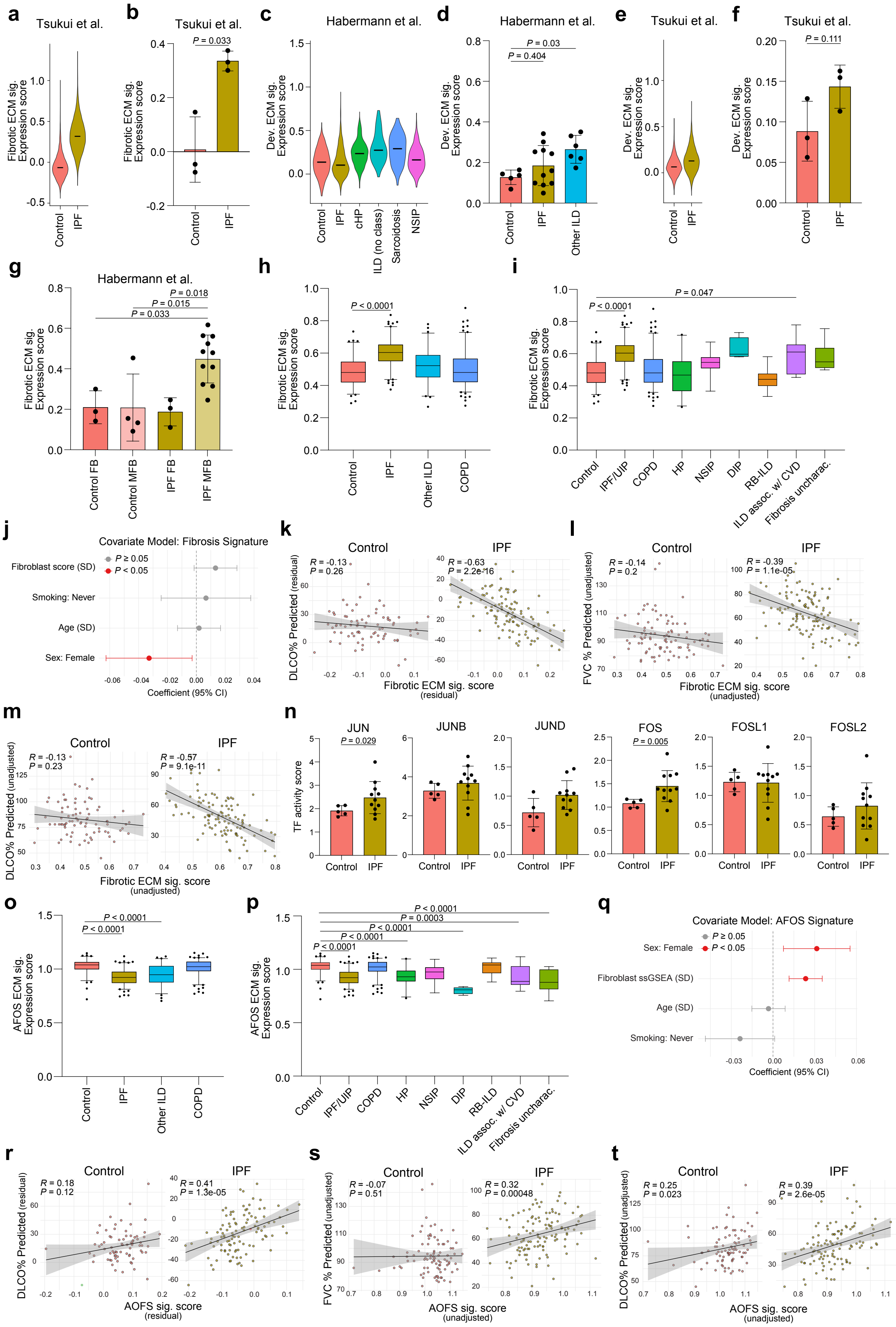

a
Tsukui et al.
Fibrotic ECM sig. Expression score
Control
IPF
b
Tsukui et al.
P = 0.033
Fibrotic ECM sig. Expression score
Control
IPF
c
Habermann et al.
Dev. ECM sig. Expression score
Control
IPF
cHP
ILD (no class)
Sarcoidosis
NSIP
d
Habermann et al.
P = 0.03
P = 0.404
Dev. ECM sig. Expression score
Control
IPF
Other ILD
e
Tsukui et al.
Dev. ECM sig. Expression score
Control
IPF
f
Tsukui et al.
P = 0.111
Dev. ECM sig. Expression score
Control
IPF
g
Habermann et al.
P = 0.018
P = 0.015
P = 0.033
Fibrotic ECM sig. Expression score
Control FB
Control MFB
IPF FB
IPF MFB
h
P < 0.0001
Fibrotic ECM sig. Expression score
Control
IPF
Other ILD
COPD
i
P = 0.047
P < 0.0001
Fibrotic ECM sig. Expression score
Control
IPF/UIP
COPD
HP
NSIP
DIP
RB-ILD
ILD assoc. w/ CVD
Fibrosis uncharac.
j
Covariate Model: Fibrosis Signature
P ≥ 0.05
P < 0.05
Fibroblast score (SD)
Smoking: Never
Age (SD)
Sex: Female
Coefficient (95% CI)
k
Control
IPF
R = -0.13
P = 0.26
R = -0.63
P = 2.2e-16
DLCO% Predicted (residual)
Fibrotic ECM sig. score (residual)
l
Control
IPF
R = -0.14
P = 0.2
R = -0.39
P = 1.1e-05
FVC % Predicted (unadjusted)
Fibrotic ECM sig. score (unadjusted)
m
Control
IPF
R = -0.13
P = 0.23
R = -0.57
P = 9.1e-11
DLCO% Predicted (unadjusted)
Fibrotic ECM sig. score (unadjusted)
n
JUN
P = 0.029
JUNB
JUND
FOS
P = 0.005
FOSL1
FOSL2
TF activity score
Control
IPF
o
P < 0.0001
P < 0.0001
AFOS ECM sig. Expression score
Control
IPF
Other ILD
COPD
p
P < 0.0001
P = 0.0003
P < 0.0001
P < 0.0001
P < 0.0001
AFOS ECM sig. Expression score
Control
IPF/UIP
COPD
HP
NSIP
DIP
RB-ILD
ILD assoc. w/ CVD
Fibrosis uncharac.
q
Covariate Model: AFOS Signature
P ≥ 0.05
P < 0.05
Sex: Female
Fibroblast ssGSEA (SD)
Age (SD)
Smoking: Never
Coefficient (95% CI)
r
Control
IPF
R = 0.18
P = 0.12
R = 0.41
P = 1.3e-05
DLCO% Predicted (residual)
AOFS sig. score (residual)
s
Control
IPF
R = -0.07
P = 0.51
R = 0.32
P = 0.00048
FVC % Predicted (unadjusted)
AOFS sig. score (unadjusted)
t
Control
IPF
R = 0.25
P = 0.023
R = 0.39
P = 2.6e-05
DLCO% Predicted (unadjusted)
AOFS sig. score (unadjusted)

**Supplementary Figure 7: Human IPF samples display elevated expression of the fibrotic ECM program and AP-1-activity.**

(a) Violin plot of expression of the fibrotic ECM program in all fibroblasts from a scRNA-seq dataset of control and IPF patients[27]. (b) Bar graph of (a) pseudobulked by patient. (c) Violin plot of expression of the developmental ECM program in all fibroblasts from a single-cell RNA-seq dataset of control patients and patients with various forms of lung disease[59]. (d) Bar graph of (c) pseudobulked by patient. (e) Violin plot of expression of the developmental ECM program in all fibroblasts from a scRNA-seq dataset of control and IPF patients[27]. (f) Bar graph of (e) pseudobulked by patient. (g) Bar graph of the control and IPF patients from (Fig. 7b) split by fibroblast type[59]. (h) Box plot of expression of the fibrotic ECM program in bulk transcriptomic data from human lung samples in control, IPF, other ILD, and COPD patients[61]. (i) Box plot of (h) where "Other ILD" samples were split into more specific disease cohorts. UIP, usual interstitial pneumonia; HP, hypersensitivity pneumonitis; DIP, desquamative interstitial pneumonia, RB-ILD, respiratory bronchiolitis-associated ILD; CVD, cardiovascular disease. (j) Forest plot of regression coefficients from a linear regression model of the fibrotic ECM program ssGSEA score against age, sex, smoking status, and fibroblast abundance score in control and IPF samples. (k) Scatter plots of the adjusted fibrotic ECM program ssGSEA score versus DLCO% predicted in (left) control and (right) IPF patients. DLCO, diffusing capacity of the lungs for carbon monoxide. (l-m) Scatter plots of the unadjusted fibrotic ECM program ssGSEA score versus (l) FVC% predicted or (m) DLCO% predicted in in (left) control and (right) IPF patients. (n) Bar graphs of expression of previously defined downstream regulons of TRE binding AP-1 family members in control and IPF fibroblasts from the dataset in (Fig. 7b)[59]. (o) Box plot of expression of the AFOS program in bulk transcriptomic data from human lung samples in (h)[61]. (p) Data from (o) where "Other ILD" samples have been split into more specific disease cohorts. (q) Forest plot of regression coefficients from a linear regression model of the AFOS program ssGSEA score against age, sex, smoking status, and fibroblast abundance score in control and IPF samples. (r) Scatter plots of the adjusted AFOS program ssGSEA score versus DLCO% predicted in (left) control and (right) IPF patients. (s-t) Scatter plots of the unadjusted AFOS program ssGSEA score versus (s) FVC% predicted or (t) DLCO% predicted in in (left) control and (right) IPF patients.

Violin plots show the median and the relative frequency of cells at a given expression value. Bar graphs show mean ± standard deviation. Box plots show the median and IQR, with whiskers extending to the 5th and 95th percentiles. Forest plots show 95% confidence intervals, and age and fibroblast abundance scores were standardized to unit standard deviation to allow comparison of effect sizes across predictors. Scatter plots show a linear regression line with 95% confidence interval. *P* values were calculated by two-tailed Welch's *t*-test (b, f, n), by ordinary one-way ANOVA with Tukey's multiple comparisons test (d, g-i, o-p; for space reasons, only significant changes from control are shown), by two-tailed t-tests of multivariable linear regression coefficients (j, q), and by Spearman's rank correlation test (k-m, r-t).
